# Repetitive extragenic palindrome (REP) elements are local, context-dependent, dual 3’UTR regulators in *Escherichia coli*

**DOI:** 10.64898/2026.03.23.713779

**Authors:** Frances E. Harris, Yanting Hu, Subhash Verma, Sankar Adhya, Weiqiang Zhou, Jie Xiao

**Author notes:** Corresponding author Jie Xiao.

## Abstract

Repetitive extragenic palindromes (REPs) are the most abundant repetitive noncoding elements in the *E. coli* genome. Despite their abundance, the primary function of REPs has remained unclear. At different times, REPs have been proposed to contribute to chromosome organization, mRNA decay regulation, and transcription termination, among other functions. Here, we show that the model REP, *REP325,* does not measurably compact the chromosome but instead acts as a 3’UTR-associated transcription regulator within the *yjdMN* operon, functioning both as a partially Rho-dependent terminator that limits transcription into the downstream *yjdN* gene and as an mRNA stabilizer that protects the upstream *yjdM* transcript from degradation. This dual role in controlling both transcriptional readthrough and susceptibility to decay provides a framework that reconciles several previously conflicting observations about REP function. Our genome-wide RNA-seq analysis further reveals that REPs with more canonical sequence and hairpin structures are more often associated with upstream-biased expression in tandem gene pairs, and that REPs positioned between convergent genes correlate with elevated expression of both genes. The large variance in expression patterns in both gene pair configurations is consistent with context-dependent termination and degradation blocking. Similarly, REPs do not uniformly affect mRNA half-lives. Because REP locations vary between *E. coli* strains, REPs likely contribute to regulatory diversity by tuning gene expression without altering protein-coding sequences or promoter regions, opening new avenues for modulating gene expression through REP-mediated transcription regulation.

## Introduction

The sparsity of noncoding DNA sequences is a defining feature of bacterial genomes. The most widely dispersed and most conserved repetitive noncoding DNA sequence motif in the *E. coli* genome is the repetitive extragenic palindrome (REP) element, which comprises ∼ 6% of noncoding DNA (1). Also referred to as Bacterial Interspersed Mosaic Elements (BIMEs), REPs have three defining characteristics: their conserved, repeated sequence motif, their exclusive location in extragenic regions, and their palindromic structure(1). The 355 annotated REPs throughout the *E. coli* genome are comprised of 1-12 palindromic units (PUs), each with one of three conserved 30-50bp sequences, termed Y, Z1, and Z2 (2, 3). REPs only occur in extragenic regions, usually within a gene’s 3’ untranslated region (UTR) (1, 4). As a consequence of their palindromic nature, REPs can form single-stranded hairpins or double-stranded cruciforms (crfDNA), both with a well-conserved mismatched bulge (**Figure 1A**). Other species of γ-proteobacteria have similar REP elements with species-specific sequences (3, 5, 6).

**Figure 1.**
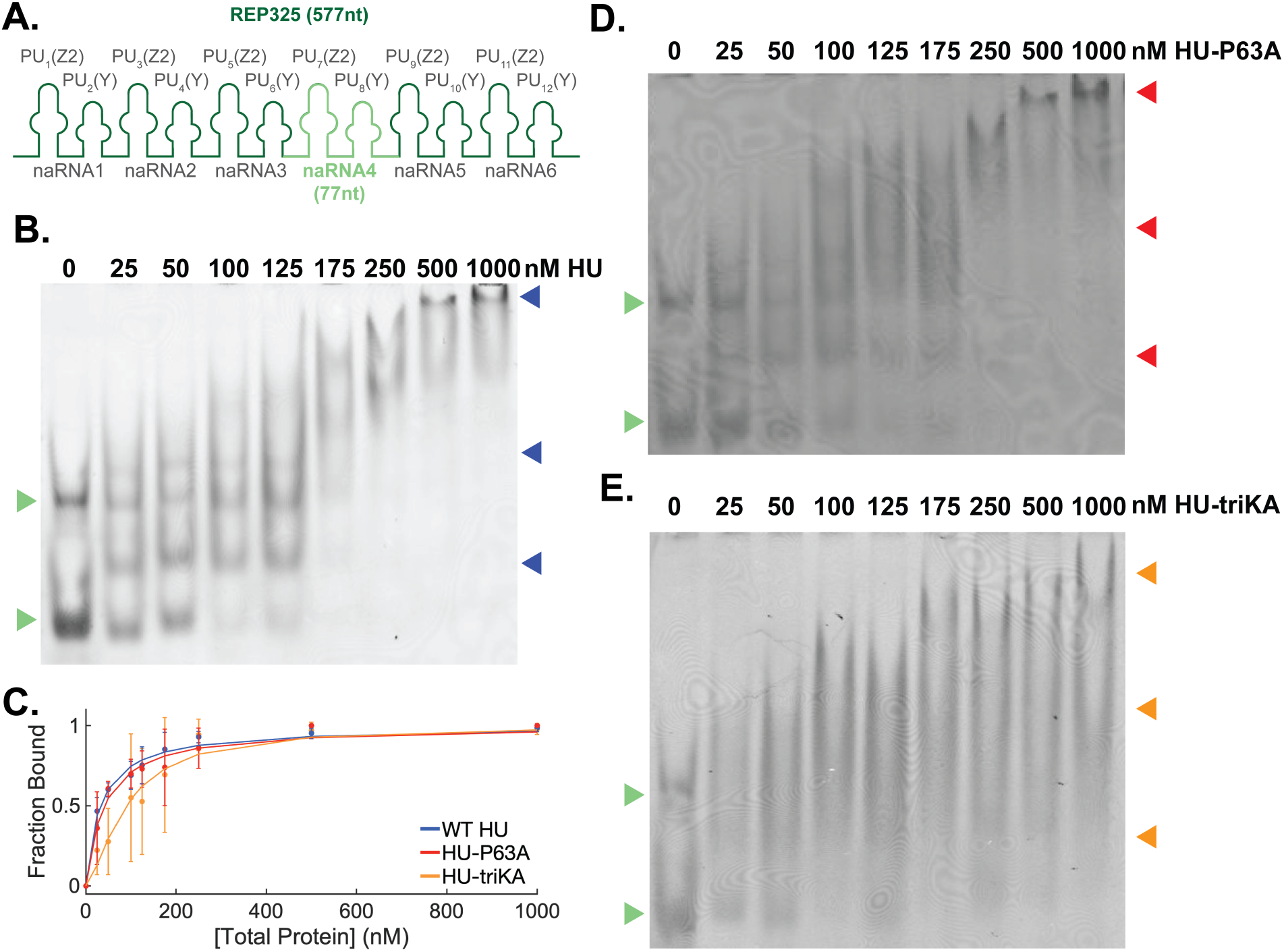
Characterization of HU binding to naRNA4. **A.** Schematics of *REP325* illustrating its 12 palindromic units (PU) and the corresponding Z2 or Y hairpins. Each two PUs are termed as one nucleoid associated RNA. naRNA4 is highlighted in light green. **B.** Electrophoresis mobility shift assay (EMSA) to probe HU-naRNA4 binding. In vitro transcribed naRNA4 (25 μM, green arrows) was incubated with increasing concentrations of purified Wildtype (WT) HUα_2_ homodimer as indicated at the top of the lanes and ran on 6% native polyacrylamide gel. HU’s binding to naRNA4 resulted two distinct, upshifted bands as indicated by the two lower blue arrows at HU concentrations ≤125 nM, and smearing, higher molecular weight bands at higher HU concentrations (top blue arrow). **C.** Binding curves of naRNA4 to HU WT, HU P63A, and HU triKA to determine the corresponding dissociation constant *K_d_*. The bound fraction of naRNA4 was calculated from EMSA gels and fit to a Hill equation. Error bars are standard deviation from the mean calculated from three replicates. WT and HU-P63A bind to naRNA with comparable affinities (*K_d_* = 32 ± 4 nM for WT HU and 41 ± 5 nM for HU-P63A), while HU-TriKA showed significantly reduced binding affinity *(K_d_* = 90 ± 9 nM). **D** and **E:** Representative EMSA gels of naRNA4 binding to HU-P63A (**D**) and HU-triKA (**E**) performed using the same condition as in B. HUP63A binds to naRNA4 similarly (red arrows) compared to WT HU, but HUTriKA showed extensive smears without distinct bands (orange arrows). Unbound naRNA4 bands are indicated by green arrows.

Given the compact and efficient nature of the *E. coli* genome, REPs have been proposed to play a structural or regulatory role since their discovery in the 1980’s. Among other potential roles (7–9), REPs have been proposed to function as chromosomal domain boundary sites, mRNA-degradation regulatory sites, and transcription-termination sites (1, 8, 10–12). These roles are not mutually exclusive and are likely context-dependent, making it difficult to delineate REPs’ primary function.

In the role of defining chromosomal domains, REPs have been proposed to connect with one another, linked by a nucleoid-associated protein HU and noncoding nucleoid-associated RNAs (naRNAs) (12–14). This model is supported by observations that naRNA4, one of the six highly structured naRNAs transcribed from a 12-hairpin REP in *E. coli REP325*, co-purified with HU (12, 13), and some REP-REP chromosomal contacts were found to be HU-dependent, *REP325*-dependent, or both (12). *In vitro*, compaction of a plasmid containing crfDNA structures required the presence of both HU and naRNA (14). However, neither the interactions within the proposed HU-crfDNA-naRNA4 complex nor the broader principles governing REP-REP contacts *in vivo* have been defined.

In the role of regulating mRNA degradation, REPs have been shown to protect upstream mRNA by physically blocking 3’→5’ exonucleases and helicases when co-transcribed in specific operons (15, 16). They also stabilized REP-containing transcripts under stress-induced ATP depletion (17). However, REP RNAs do not accumulate in wild-type (*wt*) *E. coli* cells under standard laboratory conditions (18, 19), and inserting various REPs after a reporter gene did not produce the expected stabilizing effect (20). These observations suggest that REP-mediated effects on mRNA decay may be environment- and gene context-dependent.

In the role of terminating transcription, some REPs have been identified as partial Rho-dependent terminators (RDTs), which promote RNA polymerase (RNAP) pausing and termination by the termination factor Rho (20). Consistent with other RDTs, most REPs lack the adjacent polyU track, a feature definitive of Rho-independent terminators (RITs). A few coincidental REP RITs (e.g., *REPt42*, *REPt127*, *REPt285* (4)) are the only clear exceptions. While the REP sequence motif lacks the typical C-rich and G-poor (C>G) skew in many RDTs (21), this requirement appears to be relaxed in the presence of NusG, an elongation factor and a Rho cofactor (22). Because the sequence and structural determinants of Rho-dependent termination resist clear-cut definitions, how REPs terminate transcription in different genomic and regulatory contexts remains unclear.

In this work, we investigated the roles of REPs in chromosome organization, mRNA stability, and transcription termination. We initially used *REP325* as a model REP for three reasons. First, it has already been implicated in chromosome organization. Second, *REP325* is among the longest REPs, with 12 PUs of alternating Z2 and Y sequences, making any length-dependent effects more pronounced. Third, *REP325* lies between two nonessential genes, *yjdM* and *yjdN*, so cells are likely to tolerate changes in their expression levels following genetic manipulations. Combining single-cell imaging, genetic manipulation, biochemical and transcription analyses, we discovered that the *REP325* did not appreciably contribute to nucleoid compaction as previously proposed but functioned both as a terminator and a stabilizer to modulate the *yjdMN* operon transcription in a context-dependent manner. Further genome-wide REP conservation and RNA-seq analysis revealed that while there existed large variations, REPs positioned between tandem gene pairs often favored upstream gene transcription, and that REPs positioned between convergent genes correlate with elevated transcription of both genes. Together, these analyses established REPs as local transcription modulators affecting adjacent mRNA decay and transcription termination of neighboring genes.

## Materials and Methods

### In vitro Preparation of naRNA4, crfDNA, and HU

For the binding and competition EMSAs, naRNA4 was transcribed *in vitro* using the AmpliScribe™ T7-Flash™ Transcription Kit followed by ethanol precipitation. The template was ordered from IDT (**Table S1**), and PCR amplified before use (14). Before binding reactions, RNA was refolded in 10mM Tris H-Cl, 100mM NaCl, 2mM EDTA buffer by heating to 95°C and gradually cooling to 25°C overnight with a thermocycler.

The cruciform structure was ensured by annealing 2 ssDNA oligos that have two complementary 10bp regions flanking noncomplementary hairpins (**Table S1**) (14). The oligos were ordered from IDT. An annealing reaction containing an equimolar ratio of oligos in 10mM Tris H-Cl, 100mM NaCl, 2mM EDTA was heated to 95°C and gradually cooled to 25°C overnight with a thermocycler.

HU WT and mutants were purified following the protocol used previously (23). Proteins were kept in a buffer of 50mM Tris-HCl, 150mM NaCl, and 10% Glycerol.

### Electromobility Shift Assays (EMSAs)

Binding of naRNA4 to HU and HU mutants measured by electromobility shift assay (EMSA) using native polyacrylamide gels (6% polyacrylamide 1xTBE). For binding reactions, refolded RNA (25nM) and HU (25nM-1000nM) were mixed in 1x TE buffer (final volume 5μL). The binding reactions were combined with 5μL 60% glycerol for loading. Gels were run at 4°C using 45V to load the samples and then 60V until the pink band of loading dye reached the bottom of the gel (NEB Purple Gel Loading Dye, run in empty lanes beside gel). Gels were stained with SYBR™ Gold and imaged on a Typhoon FLA9500 imager.

EMSAs were quantified using GelAnalyzer version 19.1. Lanes and peaks were manually defined. Background was subtracted using the morphological setting with a peak width tolerance of 10% of lane profile length. Bound and unbound fractions were calculated as the percentage signal in each lane based on the raw volume.

Ternary EMSAs were preformed in the same manner described above, except the gels were imaged in the Cy3 channel on the Typhoon prior to SYBR™ Gold staining. Then, the gels were re-imaged with same settings to visualize the SYBR™ Gold stain. The concentration of crfDNA (25nM) and HU (50nM) were kept constant, while naRNA4 concentration increased from 6.25nM to at least 100nM).

### Bacterial Strains and Plasmid Creation

A markerless deletion of the *REP325* element was generated using a two-step recombineering strategy. In the first step, the pBAD-ccdB-kan cassette was PCR-amplified using homology primers *REP325*-kan-ccdB-u1 and *REP325*-kan-ccdB-l1 (**Table S1**) and inserted at the *REP325* locus via homologous recombination. The pBAD-ccdB-kan cassette expresses the CcdB toxin under the arabinose-inducible pBAD promoter and confers kanamycin resistance.

Recombinant strains were selected on LB agar supplemented with 1% (w/v) glucose and 30 μg/ml kanamycin. Correct insertion of the cassette was confirmed by PCR using primers flanking the *REP*325 element and by the inability of recombinants to grow on LB agar containing 0.02% (w/v) arabinose, indicating functional expression of the ccdB toxin.

In the second step, the pBAD-ccdB-kan cassette was excised using the single-stranded oligonucleotide *REP325*-del-ss-01 (**Table S1**), synthesized by Integrated DNA Technologies. Successful markerless deletion of the *REP325* element was verified by PCR amplification followed by Sanger sequencing.

All plasmids were constructed using In-Fusion^®^ cloning (Takara Bio) and confirmed by whole-plasmid sequencing. Plasmids were transformed into the *wt* or *ΔREP325* strains by electroporation.

### DNA Staining, Nucleoid Imaging, and Nucleoid Quantification

To characterize nucleoid morphology, we imaged fixed cells stained with Hoechst dye as performed previously (24, 25). For fixation, saturated cultures were diluted 1:100 in LB with appropriate IPTG and antibiotics and grown to OD_600_ 0.4-0.6 at 37°C. Then, Hoechst 33342 dye (final concentration 10 μg/mL, bisbenzimide H33342 trihydrochloride, ThermoFisher H3570) was added to the cultures, which were incubated at 25°C for 15 more minutes with agitation. Cells were washed three times with PBS to remove excess dye and then fixed with a 3% paraformaldehyde-PBS solution at room temperature covered in foil for 15 minutes. The cells were washed two times with PBS, covered in foil, and stored at 4°C.

Cells were imaged on 3% agarose pad. Using the 3D SIM acquisition setting, 3D z-stacks of 2µm thickness (125nm slices) were collected on the GE OMX SR structured illumination microscope (excitation 405nm, emission channel 488nm, exposure 50ms, 5% of maximum laser intensity). Images were reconstructed using the SIR function with standard parameters in the GE SRx software. The maximum intensity z-projection of the 3D reconstruction was merged with the brightfield image for analysis.

A custom Matlab (v2024b) script was used to manually measure cell and nucleoid dimensions in the merged images. Fluorescence intensity set to consistent values for all images for analysis. Images from 3-4 biological replicates were analyzed blindly, resulting in 106-173 cells analyzed per condition. Since the cell dimensions were not normally distributed, the measurements were compared between conditions using a two-sided Kolmogorov-Smirnov (K-S) test.

### Hi-C Analysis

Publicly available Hi-C data was used for our analyses (Accession/SampleID- *wt* replicates: SRR7001741, SRR6354565, GSM6616791, GSM6616792. *ΔhupAB* replicates: SRR7001742, SRR6354567) (26, 27). Raw Hi-C sequencing reads were processed using HiC-Pro, which performs read alignment, quality filtering, duplicate removal, and generation of contact matrices. Contact matrices were generated at 5 kb resolution. To correct for systematic biases inherent in Hi-C data, (ICE) normalization was applied to the raw contact matrices using the built-in HiC-Pro (v.3.1.0) normalization module.

Genomic locations of REP (Repetitive Extragenic Palindromic) elements were obtained from the EcoCyc Database (4). REP loci were mapped to the Hi-C contact matrix bins at the corresponding 5 kb resolution. Binary matrices were constructed to identify bin pairs where both bins contained at least one REP element (REP–REP pairs), bin pairs where neither bin contained a REP element (non-REP pairs), and bin pairs involving bins adjacent to REP-containing bins (REP-adjacent pairs).

For each condition and replicate, contact frequencies were computed as a function of genomic distance. Specifically, for each pair of bins at a given genomic separation, the ICE-normalized contact value was extracted from the Hi-C matrix. Contact values were transformed as 1/(contact + 1) and then averaged across all bin pairs sharing the same genomic distance. The resulting distance-dependent contact profiles were smoothed using a moving average (window size = 15 bins) to reduce noise while preserving the overall trend. Contact frequency profiles were plotted separately for all loci, non-REP loci, REP-adjacent loci, and REP loci. To establish a null expectation for the contact frequency of REP loci, 100 sets of random locus pairs were generated. For each random set, the genomic distance distribution of REP–REP pairs was preserved by sampling from the observed REP pair distances without replacement. Random bin pairs were then placed uniformly across the genome at the sampled distances, and their contact frequencies were extracted from the Hi-C matrix. The minimum and maximum of the 100 smoothed random curves were displayed as a shaded region on the contact frequency plot, providing a visual reference for the expected range under the null hypothesis.

To quantify whether REP–REP and REP-adjacent contact frequencies were significantly elevated relative to random expectation, one-sided one-sample t-tests were performed at each genomic distance bin. At each bin, the 100 random contact frequency values formed the sample, and the observed REP loci (or REP-adjacent loci) contact frequency served as the test value. The alternative hypothesis was that the random values were significantly lower than the observed REP (or REP-adjacent) value, indicating elevated contact frequency at REP-associated loci. Tests were conducted both across all genomic distance bins and restricted to bins with genomic distances exceeding 0.5 Mbp. To account for multiple testing across all distance bins, p-values were corrected using the Benjamini–Hochberg false discovery rate (FDR) procedure. FDR-adjusted significance was visualized as scatter plots of −log_10_(FDR) versus genomic distance, with locally estimated scatterplot smoothing (LOESS) curves fitted for both REP loci and REP-adjacent loci, and significance thresholds marked at FDR = 0.05 and FDR = 0.01.

### Growth Curve and Drop Assays

Individual colonies were picked into 3mL of LB and appropriate antibiotic and grown at 37°C overnight. Saturated cultures were diluted 1:1000 into 500μL of LB with or without IPTG. A clear 96-well plate was loaded with two 200μL technical replicates of each dilution. The growth curve was collected by measuring the OD_600_ every 15 minutes with otherwise continuous 400rpm for 24 hours with a temperature of 37°C. The OD_600_ values were blank corrected using LB wells measured for each plate.

Each technical replicate curve was fit individually. The period beginning with the earliest occurrence of OD_600_>0.05 and ending 1.75 hours later was fit to an exponential growth model. Most cultures reached an OD_600_ of ∼0.4-0.5 by the end of the exponential growth period. The lag phase duration was defined as the time the culture took to reach an OD_600_>0.05. The sample sizes range from 3-37 biological replicates each with 2 technical replicates. The doubling time and lag phase duration parameters were compared between conditions using a two-sided t-test.

For the drop assays, saturated cultures were washed twice with 1x PBS to remove the LB media and serially diluted. 5μL of each dilution was dropped onto an LB plate with no antibiotics, and the plates were incubated at 37°C overnight. To correct for differences in saturated culture densities, the OD_600_ of saturated cultures were measured by performing a 1:10 dilution and taking the average of 3 Nanodrop OD_600_ measurements. Then, saturated culture was added to 1mL of 1xPBS such that the final OD_600_ was 0.5.

### RNA-Seq and Analysis

Individual colonies were picked into LB with appropriate antibiotic and grown to saturation overnight at 37°C. The saturated cultures were diluted 1:100 into 5mL of fresh LB with appropriate antibiotic and IPTG. The dilution was grown to exponential phase (OD_600_ 0.4-0.6) at 37°C. RNA samples were prepped as described below for Northern Blots.

Libraries were prepared using the SMARTer® Stranded Total RNA-Seq Kit v3 (Takara) and the NEBNext® rRNA Depletion Kit following the standard protocols for each. Libraries were sequenced by Azenta.

For differential expression analyses, universal Illumnia adapter sequences were trimmed using cutadapt (v.3.2) and aligned using bowtie2 (v.2.4.1) to the MG1655 reference genome (NC_000913.3). SAM files were converted to BAM using samtools (v.1.15.1). BAM files were then loaded into RStudio (v4.4.2). Gene counts were generated with featureCounts (Rsubread v.2.20.0), and differential expression analysis was preformed using DeSeq2 (v.1.46.0). Significant differential expression was determined using adjusted p-value < 0.01 and a |log_2_(fold change)|>1.5. The reads for the *yjdMN* operon were visualized using Integrated Genome Viewer (IGV_2.19.7). The y-axis was scaled by the normalization factor calculated by DeSeq2 for each replicate. For *yjdM*/*yjdN* ratio calculations, normalized counts were generated from the DESeq2 pipeline and then divided by the gene length in base pairs.

### Northern Blot RNA Sample Preparation

Individual colonies were picked into LB with appropriate antibiotic and grown to saturation overnight at 37°C. The saturated cultures were diluted 1:100 into 3-5mL of fresh LB with appropriate antibiotic and IPTG and grown to exponential phase (OD 0.4-0.6) at 37°C.

Total RNA was isolated from the cells using the Total RNA Purification from Tough-to-Lyse Samples (bacteria, yeast, plant, etc.) protocol for the Monarch Total RNA Miniprep Kit (NEB #T2010 including the optional DNAse treatment and the following two modifications:

First, to lyse the cells, the exponential culture was harvested by centrifugation (10 minutes at 4110rpm). The media was removed, and the cells were resuspended in 1mL TE. The cells were spun down using a microcentrifuge (10,000rpm for 2 minutes); the supernatant was removed. The pellet was flash frozen in dry ice and ethanol. The pellet was resuspended in 96μL 10mM Tris HCl 7.5pH 2mM EDTA, and 4μL of lysozyme (25mg/mL solution in 10mM Tris HCl 7.5pH 2mM EDTA). The cells were lysed at room temperature for 5 minutes.

Second, to increase the binding of small RNAs to the RNA purification column, twice the standard volume of ethanol was added to the flow-through before loading onto the RNA Purification column (600μL 200 proof ethanol and 300μL flow-through, loaded onto the column 450μL at a time in two spins).

The column was eluted with 100μL Nuclease-free Water. Yields usually ranged between 30-100μg. 200ng of the prep was run on an 1.0% agarose E-Gel™ to visually assess RNA quality, judged by the sharp appearance of rRNA bands. The sample was then ethanol precipitated with 1μL glycogen. The samples were resuspended in 10μL of nuclease-free water.

### Northern Blot Probe Labeling

Probes were ordered as unmodified DNA oligos from IDT (**Table S1**). The labeling reaction (final volume 10μL) contained 25pmol of unlabeled oligo, 1 μL T4 PNK enzyme (NEB), 1x T4 PNK Buffer (NEB) and 3 μL γ-^32^P. After the 30-minute incubation at 37°C, the reaction was heat inactivated (20 minutes at 65°C) and purified using the G25 Microspin column (Cytiva).

### Polyacrylamide Northern Blots

The constructs were expressed in the *wt* strain from a plasmid, preceded by a T5 promoter and followed by a lambda terminator. To keep the expressed naRNA4 and naRNA34 RNA as close to the canonical naRNA4 (77nt) and naRNA34 (177nt) as possible, both sequences were immediately followed by an rrnD terminator. Expression was induced using 1mM IPTG, and total RNA was prepared using the protocol above. For samples treated with bicyclomycin (BCM), BCM (stock concentration 12.5mg/mL H_2_O stored at −20°C, final concentration 25μg/mL) was added to the LB media at the time of dilution.

For the *in vitro* transcribed naRNA4 (IVT naRNA4) used in Northerns, a plasmid containing the template flanked by SapI and BamHI cut sites was ordered from GenScript, maxiprepped, and digested. Similar to (28), the *in vitro* transcription reaction containing 3μg DNA template, 1mM NTPs, and 2μL T7 polymerase in a final volume of 250 μL was incubated at 37 °C for 4 hours. The reaction was concentrated by ethanol precipitation and run on a denaturing gel (12.5% polyacrylamide 1xTBE 8M Urea) The desired product was purified by freeze-thawing, followed by ethanol precipitation and resuspension in nuclease-free water.

For the Northern blots, 8M Urea 1xTBE 6% polyacrylamide gels were poured with 0.75mm spacers. For each gel, samples were prepared in the smallest possible volume (5-17μL total per lane) containing 1.5-5μg of total RNA or 0.2ng-0.05ng of purified IVT naRNA4 in 1x formamide loading dye (xylene cyanol and bromophenol blue). The gel was run 15W for 5 minutes to load the samples and then 8W until the xylene cyanol dye migrated between half and two-thirds the length of the gel. If samples appeared to smile, the wattage was stepped down 2W at a time to a minimum of 2W. Gels ran for approximately 4 hours. The RNA was transferred to a nylon Amersham™ Hybond-N+ membrane by electroblotting (0.5xTBE buffer; 200mAmp; 27.5 mintues). The RNA was UV-crosslinked to the membrane (120 mJ; UV Stratalinker).

The membranes were equilibrated with Church buffer at 65°C for 20 minutes. Radiolabeled probes (25 pmol, **Table S1**) were then added to the membrane and allowed to hybridize overnight with rotation as the temperature cooled to 37°C. Dry membranes were exposed to a phosphorimager screen for 3 days.

After the exposure screens were imaged, the membranes were stripped by three 10-minute washes with boiling 0.1% SDS, reprobed with the 5S loading control probe (25 pmol, **Table S1**), re-exposed, and imaged.

For quantification, the intensity and pixel position of each band was recorded. The band intensities were quantified using ImageJ, which enforces consistent lane dimensions and vertical positions. Using relative locations of the IVT naRNA4 band (77nt) and 5S loading control band (120nt), the longest bands of the pnaRNA4 and pnaRNA34 were estimated to be the full-length transcription products containing the rrnD terminator. From the lane intensity profile, the area under a peak was used to calculate the band signal. Background subtraction was performed by drawing a straight line under the peaks. The band signal was normalized to the 5S loading control band signal.

### Agarose Northern Blots

The constructs were expressed in the *wt* strain from a plasmid, preceded by a T5 promoter and followed by a lambda terminator. Expression was induced using 1mM IPTG, and total RNA was prepared using the protocol above. The same samples were loaded twice in equal amounts on a single gel (**Fig. S11B**). Determined by the minimum RNA prep yield, 3-5μg was loaded for each sample. Since each quantification was normalized to the loading control, this difference did not affect our conclusions. RNA was transferred to the membrane by capillary wicking (5xSSC, 10 mM NaOH) for 2 hours. The RNA was UV-crosslinked to the membrane (120 mJ; UV Stratalinker). After crosslinking, the lanes were visualized by methylene blue staining and cut between the duplicate lanes. One set of lanes was probed for *yjdM* and the other for *yjdN* (25pmol, **Table S1**). Then, the membranes were equilibrated, probed, and imaged in the same manner as the polyacrylamide-based Northerns.

For quantification, band intensity and pixel position were measured in same manner as in the polyacrylamide-based Northerns. The clearest band was identified as the primary product. Product lengths were compared relative to each other, rather than using absolute lengths. Since the primary products of MR and MRN ran at the same position on the gel, the exact same boundaries were used to quantify the MR and MRN bands in each *yjdM* blot. In the *yjdN* blot, the peak widths of the endogenous *yjdN* signal in *ΔREP325* and the MN band were too different for the same constraint to be applied, but the peak locations aligned in each blot.

### Conservation Scores

In *E. coli* tandem genes, some downstream genes can be transcribed either polycistronically under the upstream gene promoter or separately under an independent promoter depending on environmental contexts or signaling factors, so we did not take current annotated promoters into account in our tandem gene definitions.

We plotted histograms for each hairpin to visualize the distribution of mapped reads at alignment positions by using Multiple Sequence Alignment (MSA v.1.36.1) package. To assess sequence conservation, we focused on positions where the mapped frequency exceeded 10% of the total possible hairpins. At each position, we calculated and plotted Shannon entropy for both the REP sequences and the random control group (randomly sampled sequences from the entire *E. coli* genome matching the REP length distribution and using identical preprocessing as with the REP sequences).

To analyze the sequence homology and construct a reference secondary structure, we extracted the dominant nucleotide at each aligned position to generate a consensus sequence and folded it using RNAfold (v2.7.0). Each REP element was then assigned a sequence conservation score defined as the mean proportion of the consensus nucleotide across its aligned positions.

To analyze the structure homology, we folded the consensus sequence by RNAfold to obtain a reference secondary structure. Each hairpin’s nucleotides were mapped to the reference positions through their shared MSA column indices. We then folded each individual hairpin by RNAfold and classified each aligned position as paired (part of a stem) or unpaired (in a loop or flanking region). Positions where a hairpin had a gap in the MSA were classified as unpaired. The structure homology score for each position was calculated as the percentage of sequences with the same paired status as the reference. For example, if the reference shows position 3 as paired but a hairpin has a gap at that position or its own fold predicts position 3 as unpaired, that hairpin is counted as non-matching at position 3. Each REP element was assigned a structure conservation score defined as the fraction of aligned positions with the same paired/unpaired status as the reference.

### Genome-Wide REP Analyses

RNA-Seq data from the *wt* condition described above was used for genome-wide REP analyses. Paired-end RNA-Seq reads were processed with UMI-aware deduplication. UMIs (14 bp) were extracted from R2 reads using umi_tools extract (v.1.1.6). Reads were then aligned to the *E. coli* MG1655 reference genome (NC_000913.3) using STAR (v.2.7.8a). PCR duplicates were removed using umi_tools dedup in paired-end mode, and BAM files were sorted and indexed with samtools (v.1.18). Read counts were then generated for each defined region across all replicates using GenomicAlignments (v.1.40.0) in R (v.4.4.0), counting fragments falling entirely within each region. Read counts were normalized by library size and averaged across biological replicates.

Random control gene pairs were constructed from adjacent tandem or convergent gene pairs that lacked REP elements. Gene pairs were filtered to match the intergenic distance range observed among REP-associated pairs (less than or equal to the maximum REP intergenic distance), and those with zero or negative intergenic distances were removed. This filtering yielded 2,451 candidate tandem pairs and 383 candidate convergent pairs. For statistical comparisons, size-matched random subsets were drawn (n = 146 for tandem pairs, n = 207 for convergent pairs) to match the number of REP-associated gene pairs.

Relevant expression patterns were compared between REP-associated and control pairs using two-sided Wilcoxon rank-sum tests. Based on our *REP325* results, we used the log2((upstream + 1) / (downstream + 1)) ratio for tandem gene pairs to quantify the relative expression difference, where a pseudo count of 1 was added to both numerator and denominator to stabilize the calculation when expression values were low. For convergent gene pairs, we used log2(expression + 1) values. To examine the relationship between REP characteristics and gene expression, we stratified the analysis by sequence and structure conservation scores (divided into quartiles based on conservation analysis).

### Half-Life Analysis

To investigate whether REP elements influence mRNA stability, we analyzed mRNA half-life data from Esquerré *et al.*(29), using their 0.1^−1^hr growth rate condition. Because gene expression levels differed significantly between REP-associated genes and the random control group, we constructed a set of expression-matched controls to account for this potential confounding factor. For each REP-associated gene, we identified a matched random control gene whose average expression level fell within ±1% of the REP-associated gene’s expression. Matching was performed separately for tandem and convergent configurations. REP-associated genes without a suitable match were excluded from analysis. Half-life values were compared between REP-associated genes and expression-matched control genes (n = 104 for tandem genes, n = 210 for convergent genes) using two-sided Wilcoxon rank-sum tests.

## Results

### HU binds naRNA4 with moderate affinity

Previous work proposed that the 12-hairpin long *REP*325 is transcribed and processed into six two-hairpin segments, termed naRNAs, each containing a Z2 and a Y PU (**Fig. 1A**). One of them, naRNA4, was shown to complex with HU and crfDNA driving chromosomal compaction (12–14). However, the presence of the HU-crfDNA-naRNA4 complex has not been thoroughly established. To address this gap, we characterized the interactions between HU, naRNA4, and crfDNA using an electromobility shift assay (EMSA).

We incubated purified HU (HUα_2_ homodimer) at varying concentrations with a fixed amount of *in vitro* transcribed naRNA4 and analyzed complex formation on a 6% native polyacrylamide gel stained with SYBR™ Gold (Methods). As in previous work, the *in vitro* transcription products of naRNA4 ran as two bands of different lengths on the native gel (**Fig. 1B, Fig. S1A-B**, green arrows) (12). At HU concentrations below 125 nM, we observed distinct bands of HU-naRNA4 (**Fig. 1B, Fig. S1A**, two lower blue arrows), similar to previously observed binding of HU to a small, structured RNA dsrA at sub-100 nM range, indicating well-defined binding stoichiometry(30). At intermediate HU concentrations (175-225 nM), we observed a smearing pattern at higher molecular weights, which diverged from multiple discrete bands at increasingly higher molecular weights observed for HU’s binding to crfDNA(23), indicative of larger but weakly associated complexes that disassociate during electrophoresis. At 500 nM and above, HU-naRNA4 formed a saturated, higher ordered oligomeric complex (**Fig. 1B, Fig. S1A**, top blue arrow). In contrast, HU’s binding to unstructured polyU RNA produced a single, faster-migrating band in the μM range (**Fig. S1C-E**). While EMSA bands cannot be interpreted as absolute stoichiometric ratios, these results suggested that HU binds structured naRNA4 with higher affinities than nonstructured polyU and has a better-defined stoichiometry at low HU concentrations than at higher HU concentrations.

To quantify the binding affinity between HU and naRNA4, we measured the intensity of the two bands corresponding to free naRNA4 at each HU concentration and assigned the remaining lane intensity to HU-naRNA4 complexes. Fitting the data to a Hill equation yielded an apparent K_d_ of 32 ± 4 nM for HU-naRNA4 (μ ± s.d., n = 3 repeats, **Fig. 1C, Fig. S1F, Table S2, Table S3**), similar to the reported K_d_ of 22 nM for HU-dsrA binding (30). This affinity is slightly weaker than HU-crfDNA binding (Kd ∼ 5 −25 nM (31, 32)) but significantly tighter than HU binding to linear dsDNA, which ranges from 200 nM to 2.5 µM depending on the ionic conditions(23), or to polyU RNA, which we measured in the µM range (K_d_ = 4.3 ± 0.1 µM, **Fig. S1D-E**).

### HU mutants alter crfDNA and naRNA4 binding differentially

Previous studies have shown that HU engages DNA through two distinct binding modes: a structure-specific mode that relies on P63-mediated base intercalation and a non-specific mode that depends primarily on surface lysine (K3, K18, and K83)-mediated electrostatic interactions (**Fig. S1G** (23, 25, 33, 34)). However, the effect of these mutations on HU-naRNA binding is unknown (30, 35). Using the same EMSA conditions as with WT HU, we measured the binding of HU-P63A and HU-triKA (K3A/K18A/K83A) to naRNA4. HU-P63A displayed a naRNA4 binding pattern qualitatively similar to WT HU, with one discrete band at low HU concentrations, smeared bands at intermediate concentrations, and a high mobility oligomeric complex at saturating HU concentrations (**Fig. 1D, Fig. S1H)**. The apparent K_d_ of HU-P63A-naRNA4, 41 ± 5 nM (n =3 replicates) was comparable to that of WT HU, indicating that proline intercalation is not critical for HU-naRNA4 binding **(Fig. 1E, Fig. S1F)**. In contrast, HU-triKA binding to naRNA4 produced no discrete bands at any HU concentration and instead yielded only smeared signals characteristic of weak, heterogenous association (**Fig. 1E, Fig. S1I**). The apparent K_d_ of HU-triKA-naRNA4 binding was 90 ± 9 nM (n = 3 replicates, **Fig. 1C, Fig. S1F**), significantly weaker than WT HU-naRNA4 binding. Together, these results indicate that HU binds naRNA4 with high affinity, but the binding does not appear to rely on the proline intercalation mechanism observed in HU-DNA cocrystal structures (36).

### HU-crfDNA-naRNA4 does not form a stable ternary complex

Next, we designed a ternary complex EMSA to probe the potential formation of a defined complex between HU, crfDNA and naRNA. (**Fig. S2**). We constructed a synthetic cruciform crfDNA structure by annealing two ssDNA oligos with complementary 10bp arms but noncomplementary hairpin regions to form a synthetic cruciform structure (**Table S1, Methods)** as described previously (14). One strand of the crfDNA was end labeled with a fluorescent dye Cy3. We incubated 50 nM HU, 25 nM Cy3-labeled crfDNA with an increasing concentration of naRNA4 from 6.25 nM to 100-200 nM at 37°C for 45 min and then ran the mixture on a native polyacrylamide gel. These concentration ranges are similar to the concentrations used in AFM previously (5nM DNA, 50nM naRNA4, and 100nM HU (14)). At 50 nM HU, we observed apparent complex formation between HU and crfDNA (**Fig. S2, lane 2**). At increasing naRNA4 concentrations, we did not observe a distinct band migrated at a different molecular weight where both naRNA4 (visualized by SYBR™ Gold) and crfDNA (visualized by Cy3 fluorescence) were present; the crfDNA banding pattern in the Cy3 channel mimicked that in the HU-crfDNA only condition regardless of the naRNA4 amount (**Fig. S2**). In addition, we observed a slight increase in unbound crfDNA at higher naRNA4 concentrations, suggesting that naRNA4 competed out crfDNA for HU binding. We concluded that HU does not form a ternary complex with crfDNA and naRNA4 under our experimental conditions.

### Deletion of REP325 does not affect nucleoid volume

Although the ternary EMSA as described above did not reveal a stable HU-naRNA-crfDNA ternary complex *in vitro*, our results do not exclude the possibility of *in vivo* complex formation and chromosomal compaction as previously proposed (12, 14). To assess the proposed nucleoid compaction role of *REP*325 quantitatively, we constructed a marker-less *ΔREP325* strain (Strain *ΔREP325*, **Table S1**) from a previous *REP*325 deletion strain in the MG1655 background (13) by removing the *cat-sacB* cassette that had replaced the *REP*325 element (577 bp) between the upstream *yjdM* and downstream *yjdN* genes on the chromosomal locus (Materials and Methods). These *ΔREP325* cells had indistinguishable cell size distributions compared to wildtype (*wt*) parental cells (**Fig. S3A**, **Table S4, Table S5**), We grew *wt* and *ΔREP325* cells to exponential phases (*OD_600_* ≈ 0.4 - 0.6) in LB medium at 37°C, stained cells with DNA-intercalating Hoechst dye, fixed the cells, and imaged nucleoids by structured illumination microscopy (SIM) (**Fig. 2A**) as we previously described (24, 25). The sub-diffraction-limited resolution of SIM (∼ 100 nm) (37) allows for accurate measurement of nucleoid dimensions. As shown in **Fig. 2B**, in both strains, the nucleoid occupied ∼ 60% of the cell area and the nucleoid area distributions did not differ significantly from each other (**Table S5,** *p*-value: 0.36). Our results demonstrated that the loss of *REP*325 did not appreciably affect nucleoid volume as previously suggested.

**Figure 2.**
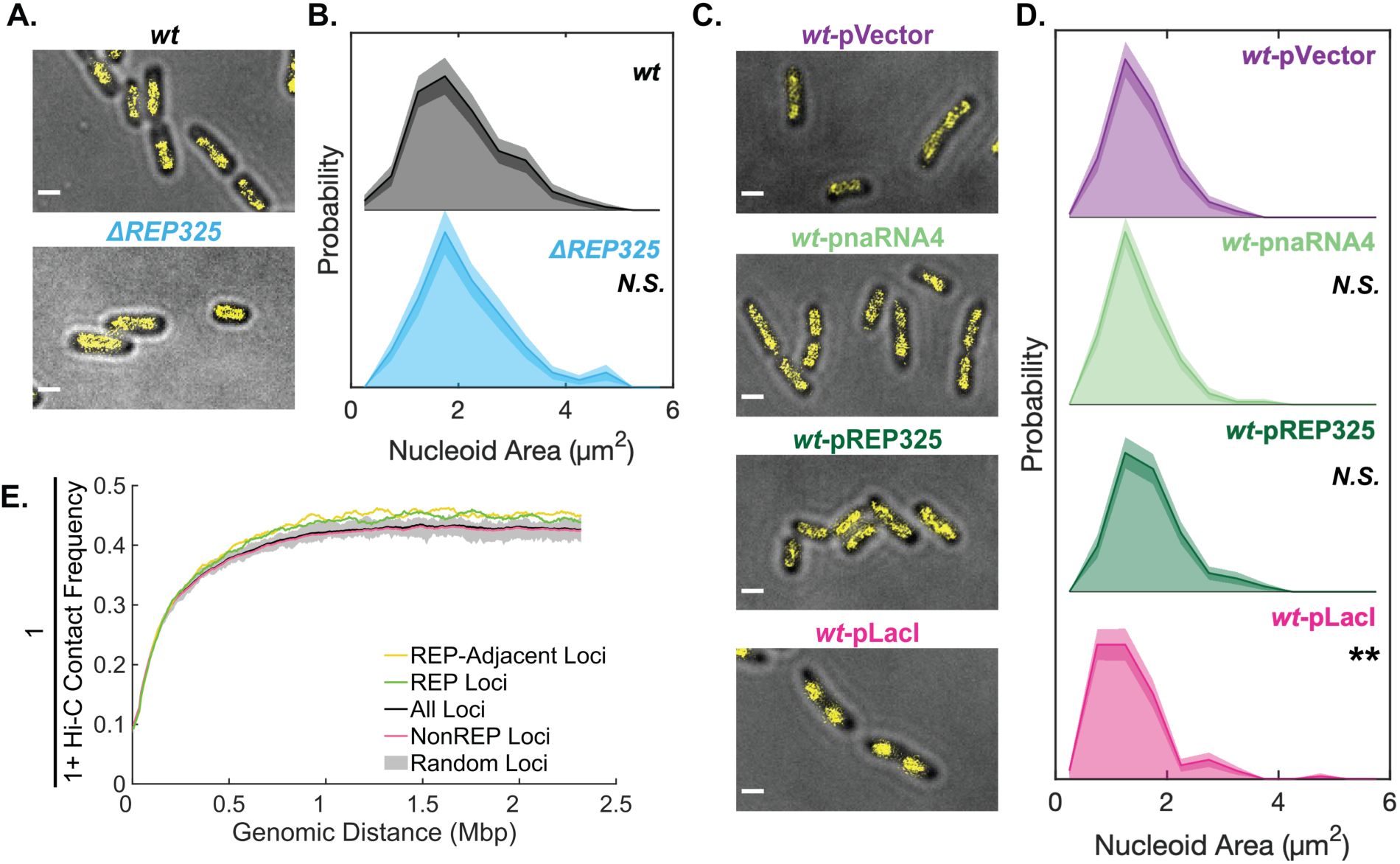
*REP325* does not affect nucleoid compaction. **A.** Representative SIM (structured illumination microscopy) images of *wt* and *ΔREP325* cells (gray) with nucleoids stained with the Hoechst dye (yellow). Scale bar: 1 μm. **B.** Nucleoid area distributions calculated from SIM images showed no significant difference between *wt* (gray) and *ΔREP325* (cyan) cells. Significance determined by *p*-value <0.05 via two-sided KS-test. **C.** Representative SIM images of Hoechst dye-stained *wt* cells harboring pVector (1 mM IPTG), pnaRNA4(1 mM IPTG), pREP325 (1 mM IPTG), or pLacI (0 mM IPTG). Scale bar: 1μm. **D.** Corresponding nucleoid area distributions of the four strains showed that only cells harboring pLacI (pink) had significant compaction compared to the pVector control (lavender). Significance determined by *p*-value <0.05 via two-sided KS-test **E.** REP-REP contact frequency analysis of a Hi-C dataset (27) showed that the apparent separations between REP-REP loci (green) or REP-adjacent loci (yellow) were slightly larger than those of all chromosomal loci (black), NonREP loci (pink), or random loci (gray) at the same genomic distances. The apparent separation (y axis) was calculated using the inverse average contact frequency at each genomic distance with moving average across 15 bins (bin size, 5kb).

### Overexpression of REP325 or individual naRNAs did not affect nucleoid volume

We reasoned that the negligible impact of *ΔREP325* on the nucleoid volume might reflect low naRNA4 transcription from the *wt* chromosomal locus. To examine whether elevated transcription of naRNA or REP325 could promote nucleoid compaction, we cloned *naRNA4* or *REP325* under an IPTG-inducible *P_T5_* promoter onto a mid-copy number plasmid (Strains *wt*-pnaRNA4 and *wt*-pREP325 respectively, **Table S1**) and compared the nucleoid volumes of the two strains to a *wt* strain carrying an empty vector (*wt*-pVector, **Table S1**). *Wt*-pnaRNA4 cells were marginally smaller than the *wt*-pVector cells under the same growth and induction conditions (**Fig. 2C, Fig. S3D**, **Table S5**), but their nucleoid areas were not significantly different (**Fig. S2C-D, Table S5**). Similarly, overexpressing the entire *REP325* element (*wt*-pREP325) from the same *P_T5_* promoter on the plasmid did not cause any appreciable changes in the nucleoid area (**Fig. 2D**, dark green, **Table S5**), although *wt*-pREP325 cells showed a modest but statistically significant increase in cell area (**Fig. S3D, Table S5**). The cell area difference for *wt-*pREP325 cells seemed attributable to differences in cell width rather than length (**Fig. S3E-F, Table S5**). Thus, in the *wt* background, neither naRNA4 nor the full-length REP325 overexpression affected the nucleoid volume significantly.

Because previous work reported nucleoid compaction upon naRNA4 or REP325 expression in a *ΔREP325* background (12, 13), we considered the possibility that the *ΔREP325* background might be particularly sensitive to REP325- or naRNA4-driven compaction. We therefore measured the nucleoid dimensions of cells overexpressing naRNA4 or REP325 in the *ΔREP325* background. However, nucleoid area distributions for *ΔREP325*-pREP325 and *ΔREP325*-pnaRNA4 cells did not differ significantly from the *ΔREP325*-pVector control (**Fig. S4**, **Table S5**). These results indicated that increased expression of neither naRNA4 alone nor full-length REP325 produces detectable nucleoid compaction under our conditions, in either *wt* or *ΔREP325* backgrounds.

### Overexpression of LacI significantly compacts the nucleoid

To verify our imaging and analysis pipeline were sensitive enough to detect changes in nucleoid volume, we overexpressed the *lac* repressor LacI from a plasmid (Strain *wt*-pLacI, **Table S1**), which previously was shown to compact the nucleoid due to nonspecific DNA binding (38). Under the same growth and imaging conditions, *wt*-pLacI cells displayed significantly condensed nucleoids compared to *wt*-pVector cells (**Fig. 2D**, bottom row), and the nucleoid area distribution shifted markedly toward smaller values (**Fig. 2D**, pink**, Table S5**). The highly condensed nucleoid was accompanied by chromosome segregation defects: ∼ 8% (n = 148 cells) showed asymmetric nucleoids or appeared anucleate. In contrast, such chromosome segregation defects were essentially absent in *wt*-pnaRNA4 (0 out of 146) or *wt*-pREP325 (1 out of 146) cells, further supporting the conclusion that the overexpression of individual naRNAs or full-length REP325 did not appreciably perturb the nucleoid morphology.

### REP-REP pairs do not form long-range chromosomal contacts in the E. coli chromosome

If REPs served as points of contact between distant genomic loci to compact the chromosome as previously proposed (8, 12), chromosomal loci containing REP*s* should exhibit elevated long-range interactions compare to loci of similar genomic distances. To test this possibility, we analyzed previously published *E. coli* Hi-C data sets (26, 27) and computed contact frequencies as a function of genomic distances for pairs of genomic bins containing annotated REPs (REP loci, 310/928 bins, **Fig. 2E**, green, **Fig. S5A**). For comparison, we also computed contact frequencies for bin pairs not containing REPs (nonREP loci, 618/928 bins, **Fig. 2E**, pink, **Fig. S5A**), for all loci (**Fig. 2E**, black, **Fig. S5A**), and for 100 sets of 355 random chromosomal loci, each sampled to match REP loci’s pairwise genomic distance distribution (Random loci, **Fig. 2E**, gray, **Fig. S5A**). To control for possible mapping biases of repetitive REP sequences in the Hi-C data, we also quantified contacts between bins directly adjacent to REP loci (REP-adjacent loci, **Fig. 2E**, yellow, **Fig. S5A**). As shown in **Fig. 2E** and **S5A**, REP or REP-adjacent loci did not show enhanced long-range contact frequencies; instead, they exhibited slightly lower contact, or were spaced further apart, at the same genomic distances than nonREP loci, all loci, and random loci. Note that because nonREP loci constituted the majority of genomic loci, they dominated the genome-wide average contact probability. In summary, we concluded that REPs do not form compacting long-range chromosomal contacts.

### REP-REP contacts are not dependent on HU

Next, to further examine whether the observed REP-REP contacts were dependent on HU, which is known to promote long-range DNA-DNA interactions in *E. coli* (27), we analyzed published *E. coli* Hi-C data collected from an HUαβ deletion strain (*ΔhupAB*, (27)). As noted in the original publication, the *ΔhupAB* strain showed a global reduction in long-range contacts compared to *wt*, evidenced by a higher plateau in the contact–distance curve beyond ∼0.5 Mb for all loci (**Fig. S5B**). In *ΔhupAB*, REP loci and REP-adjacent loci had consistently lower contact frequencies than other loci, with an effect size similar to that seen in *wt* given the overall reduction in long-range contacts. Thus, although HU generally facilitates long-range chromosomal contacts, loss of HU does not preferentially disrupt REP-REP interactions, suggesting that REP-REP interactions are not dependent on HU.

We also considered the possibility that a few REPs, instead of all REPs, might uniquely form long-range contacts, as previously reported for several specific REP-REP pairs in 3C experiments (12). We examined the Hi-C contacts of these specific REP-REP pairs and found that some, but not all pairs, had higher contact frequencies relative to the genomic average (**Fig. S5C**). In addition, the Hi-C data did not recapitulate the reported HU-dependence of these specific pairs. For example, some pairs previously reported as HU-dependent exhibited higher contact frequencies in *ΔhupAB* than in *wt* Hi-C maps (**Fig. S5D**). Therefore, we conclude this specific subset of REPs is not organizing the chromosome.

Taken together, our analysis of Hi-C data suggested that REPs do not drive global, universal long-range chromosomal contacts. While the effect was not strong enough to claim that the REP-containing loci were actively repelled from each other, the data did not support a model in which widespread long-range REP-REP contacts, HU-dependent or otherwise, appreciably contribute to chromosomal compaction.

### Deletion of REP325 accelerates cell growth

Our results so far showed that neither deletion nor overexpression of *REP325* altered nucleoid compaction and that REP-REP contacts are an unlikely mechanism of chromosome organization. Interestingly, when we measured the growth rates of *ΔREP325* and *wt* cells using a plate reader-based assay, we observed that *ΔREP325* cells grew faster (29 ± 2 minutes, μ ± std for all growth parameters, N = 37 biological replicates, **Table S6, Table S7**) and had a shorter lag phase (2.1 ± 0.4 hours, **Fig. S6**) compared to *wt* cells (35 ± 2 minutes and 2.3 ± 0.4 hours for doubling time and lag phase duration respectively, N = 30 biological replicates, **Table S7**). Importantly, our *wt* strain did not differ from the *E. coli* Genetic Resource Center *wt* (*N* = 3 biological replicates, **Table S7**), indicating that background *wt* mutations did not bias our growth measurements. The *ΔREP325* differences were modest but statistically significant (*p* < 0.05, two-tailed *t*-test for all growth parameter comparisons, **Table S7**).

To test whether these growth effects were due to changes in REP RNA expression, we measured the growth of *ΔREP325* cells carrying plasmids overexpressing the entire REP325 element (pREP325) or naRNA4 only (pnaRNA4) **(Fig. S7)**. Neither plasmid significantly altered the growth rate or lag phase duration in the *ΔREP325* background (**Fig. S7C-D, Table S7**). Similarly, overexpressing REP325 or naRNA4 in the *wt* background did not appreciably affect growth (**Fig. S8D, Table S7**). In contrast, overexpression of LacI from the same plasmid significantly slowed growth (**Fig. S8D, Table S7**), an expected consequence of an over-compacted nucleoid (38). Because neither REP325 nor naRNA4 expression restored *wt*-like growth in *ΔREP325* cells, we concluded that the increased growth rate of *ΔREP325* cells arises from a mechanism other than reduced REP RNA expression, further confirming that *REP325* does not compact the nucleoid as previously proposed.

### Upstream yjdM knockout phenocopies the accelerated growth of ΔREP325 cells

Since *REP325* RNA expression itself did not affect cell growth, we reasoned that *REP325* deletion might instead affect expression of its neighboring genes (1, 10, 11). On the chromosome, *REP325* is flanked by *yjdM* upstream and *yjdN* downstream (**Fig. S6A**). Both genes are nonessential and poorly characterized. The *yjdMN* operon lies immediately upstream of the phosphonate metabolism operon *phnC-P* but is not required for phosphonate utilization (39), yet purified YjdM protein was recently shown to catalyze a phosphonate-related reaction *in vitro* (40). As a possible consequence of *REP325* deletion (1, 10, 11), altered *yjdM* and/or *yjdN* expression could impact cell growth by affecting metabolism.

To test whether altered *yjdM* expression affects growth, we examined the growth of a *yjdM* knockout strain from the Kieo collection (41) (*ΔyjdM*, **Table S1**). We found that *ΔyjdM* cells grew similarly to *ΔREP325* cells with a shorter lag-phase duration of 2.2 ± 0.1 hours and a faster doubling time of 30 ± 1 minutes compared to *wt* (**Fig. S6**, **Table S7**, *p*-values > 0.05). These results suggested that the altered growth phenotype of *ΔREP325* cells was likely caused by diminished expression of the *yjdM* gene.

### Upstream yjdM overexpression prolongs lag phase

To further test whether the altered growth phenotype was related to *yjdM* expression, we overexpressed the *yjdM* gene from a plasmid (pM) in our *wt* background (**Fig. S9B**). Compared to the vector-only control, *yjdM-*overexpressing cells exhibited a markedly prolonged lag phase (6 ± 1 hr, **Fig. S9C**, blue) but a similar doubling time once the exponential growth phase began (38 ± 2 min, **Fig. S9D, Table S6, Table S7**). In contrast, overexpressing the downstream *yjdN* gene (pN) produced growth curves indistinguishable from the vector-only control (**Fig. S9B-E**, red curve). These results demonstrated that the under- or overexpression of the upstream *yjdM* gene, but not the downstream *yjdN* gene, was associated with altered growth phenotypes. They also suggested that *REP325* deletion may destabilize *yjdM* mRNA, phenocopying *yjdM* knockout.

To further investigate how *REP325* in the operon affects the expression of the neighboring *yjdM* and *yjdN* genes, we construct plasmids expressing different *yjdMN* operon combinations, including *yjdM*-*REP325* (pMR), *yjdM*-*REP325*-*yjdN* (pMRN), and *yjdM*-*yjdN* (pMN, **Table S1**), and measured their growth phenotypes (**Fig. S9A-B**). Cells expressing *yjdM*-*REP325* (pMR) or the full operon (pMRN) exhibited varied and extended lag phases (**Fig. S9D**, light blue and yellow curves, **Table S7**), comparable to cells expressing *yjdM* only (pM). However, cells expressing *yjdM*-*yjdN* without REP325 (pMN) interestingly had a lag phase indistinguishable from that of the vector control. Furthermore, all constructs showed doubling times similar to that of the vector-only control (**Fig. S9D, Table S7**).

Extended lag phases are often associated with cell stress or reduced viability in stationary phase cultures used as inocula (42, 43). To compare the number of viable cells in stationary phase cultures of these constructs, we used a serial dilution colony formation assay (Methods). We observed that pM, pMR and pMR cells showed lower fractions of viable stationary phase cells (**Fig. S9E**) and lower stationary phase culture densities (**Fig. S9F**), consistent with their reduced viability in stationary phase cultures. Although the underlying biology of this reduced viability remains unclear, the effect was reproducible and supported a model in which dysregulation of *yjdM* expression, modulated by *REP325*, underlies the observed growth phenotypes.

### REP325 modulates transcription within its operon

Our results above showed that the altered growth phenotypes were reproducible and likely caused by dysregulation of *yjdM* expression, modulated by *REP325*. To investigate directly how *REP325* shapes the expression of the *yjdMN* operon, and how this effect might in turn alter the global transcriptional landscape and consequently the physiological state, we performed RNA-seq on *wt* and *ΔREP325* cells, as well as in cells overexpressing *yjdM* (*wt*-pM).

We first examined the effect of *REP325* deletion on *yjdM* and *yjdN* transcription in the *yjdMN* operon (**Fig. 3A**). In *wt* cells, the upstream *yjdM* gene was transcribed at a significantly higher level than the downstream *yjdN* gene (normalized mRNA counts *yjdM*/*yjdN* = 35.3±14.3). In *ΔREP325* cells, however, the *yjdM*/*yjdN* expression ratio markedly decreased to 1.02±1.02. These ratio changes were also reflected in the differential expression analysis (**Fig. S10A, Table S8**), where *yjdM* transcription decreased (log_2_FC = −1.74, *p*_adj_ = 6.2×10^−13^) whereas *yjdN* transcription increased (log_2_FC = 3.36, *p*_adj_ = 1.8×10^−10^) in *ΔREP325* when compared to *wt* cells. The reciprocal changes in *yjdM* and *yjdN* expression in the absence of *REP325* are consistent with *REP325* having a dual role regulating mRNA stability and termination in the operon.

**Figure 3.**
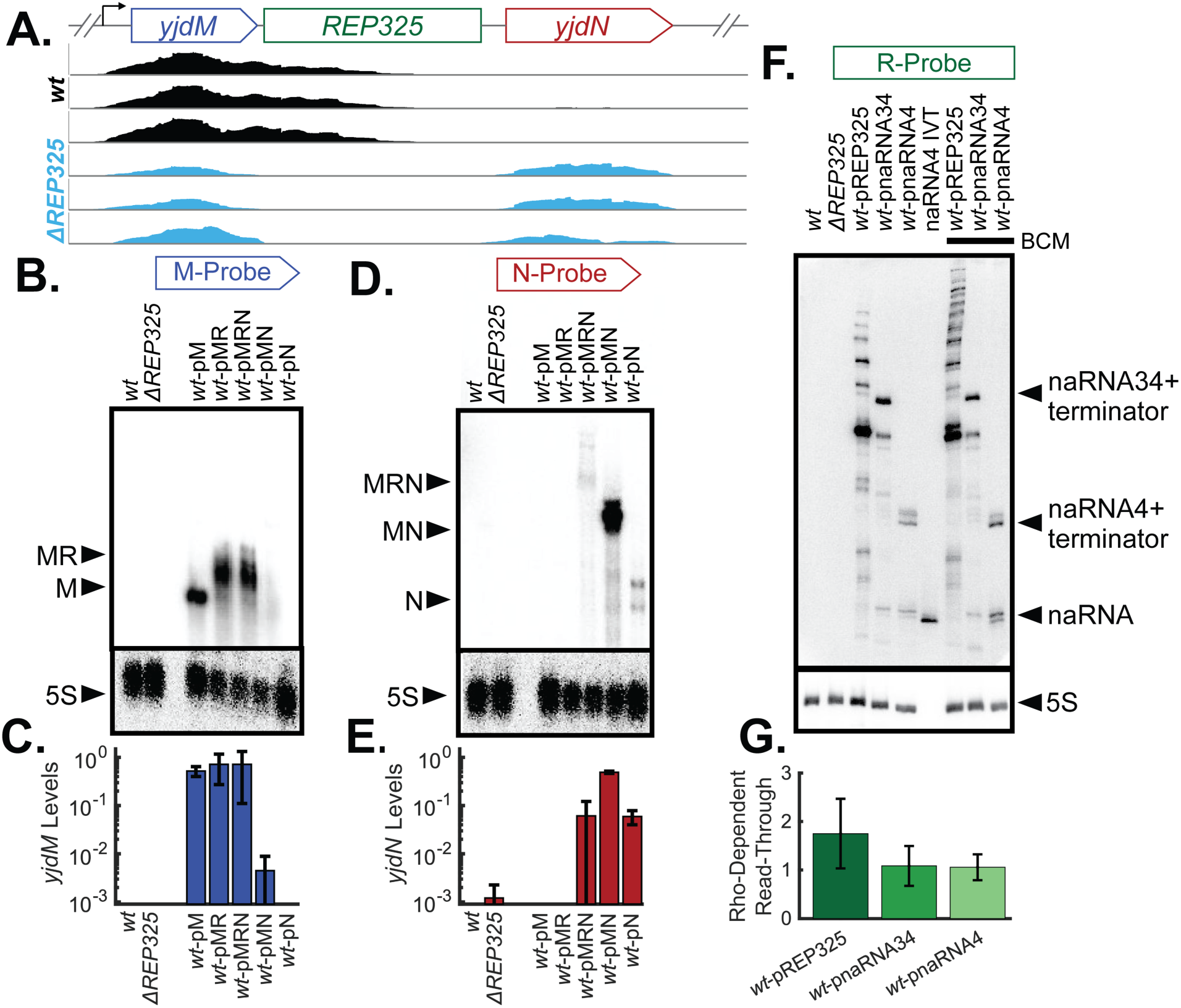
*REP325* stabilizes *yjdM* in *yjdMN* operon and acts as a partial Rho-dependent terminator. **A.** Normalized RNA-Seq read coverage (three repeats) in the *yjdMN* region (NC_000913.3:4,326,900-4,325,100) showed reduced *yjdM* transcription and enhanced *yjdN* transcription in *ΔREP325* cells (cyan) compared to wt cells (black). **B.** A representative Northern blot of *yjdM* transcription probed using the M-probe in *wt*, *ΔREP325*, and *yjdMN* operon overexpression plasmids conditions (pM, pMR, pMRN, pMN, and pN). The primary bands of *yjdM* transcript (M) and *yjdM* plus partial *REP325* (MR) were indicated by arrow heads on the left. Total cellular 5S RNA (bottom) was run as a loading control for all Northern gels. **C.** Corresponding quantifications of *yjdM* expression levels from B. showed negligible yjdM transcription in wt cells, but robust transcription in cells harboring plasmid constructs pM, pMR, and pMRN, but significantly reduced transcription in cells harboring pMN, indicating the stabilization of *yjdM* transcript by REP325 only in the presence of *yjdN*. **D.** A representative Northern blot of *yjdN* transcription using the N-probe for the same samples as described in B. The primary bands of *yjdN* transcript (N), *yjdM* -*yjdN* (MN), and the full-length operon *yjdM-REP325-yjdN* (MRN) were indicated by arrow heads on the left. **E.** Corresponding quantifications of *yjdN* expression levels from C. showed enhanced transcription of *yjdN* in *ΔREP325* cells compared to *wt* cells, and in cells harboring plasmid pMN (*yjdM-yjdN* without REP 325) compared to cells harboring the full-length operon (pMRN) or just *yjdN* (pN). **F.** A representative Northern blot of REP325 transcription using the R-probe in *wt, ΔREP325,* and *wt* cells harboring plasmid constructs pREP325, pnaRNA34, and pnaRNA4 in the absence and presence of Rho-inhibitor BCM. Expected sizes of transcription products of naRNAs were shown by arrowheads on the right. **G.** Quantified Rho-dependent read-through for pREP325, pnaRNA34, and pnaRNA4 based on the Northen blot in F. For all quantifications, error bars are standard deviations of the mean.

Next, we compared the global transcriptional profiles of *ΔREP325* and *wt*-pM overexpression cells to *wt* cells to investigate whether perturbing *REP325* or *yjdM* expression produced global transcriptional changes indicative of a generalized stress response. As shown in **Fig. S10A**, *ΔREP325* cells showed a modest number of differentially expressed genes (99 genes downregulated, 59 genes upregulated, abs(log_2_FC)>1.5 and *p*_adj_<0.01). Upregulated genes showed no significant GO term enrichment (**Table S9**), no coherent enrichment of classic stress-response programs, and spanned diverse functional categories including metabolism and acid stress response related genes (*gutM*, *appY*, *gadY*) and several “y-genes” of unknown function. Downregulated genes included mostly flagellar genes (**Table S9**), a pattern previously reported as an *E. coli* stress response (44, 45). We also tested whether these *ΔREP325* differentially expressed (DE) genes were related to changes in global chromosomal organization by examining if they were HU-regulated, *REP*-adjacent, or supercoiling-sensitive **(Table S11**, Supplementary Info), as has been previously proposed (12). However, we concluded that deleting *REP325* did not affect gene expression through global chromosomal structuring mechanisms.

In cells overexpressing *yjdM*, we observed relatively few differentially expressed genes (37 genes upregulated, 45 genes downregulated, abs(log_2_FC)>1.5 and *p*_adj_<0.01, **Table S12**). *Wt*-pM cells had no clear transcriptional signatures of stress responses, and enriched GO terms were limited to processes such as mechanosensory behavior, magnesium ion responses, and hexose catabolic process, none of which relates to YjdM’s proposed role in phosphonoacetate metabolism. Taken together, we concluded that deleting *REP325* affected *yjdM* and *yjdN* transcription, but under or overexpressing YjdM did not trigger a recognizable global stress response. Further studies of YjdM function specifically may help link *yjdM* dysregulation to the observed growth differences.

### REP325 stabilizes yjdM mRNA in the yjdMN operon

The RNA-seq experiment of *ΔREP325* cells showed clear modulation of *yjdM* and *yjdN* transcription by *REP325*, but typical RNA-Seq read length (∼150bp) prohibits detection of intact *yjdM*-*REP325* or *yjdM*-*REP325-yjdN* mRNA to directly probe the role of REP325 in modulating *yjdM* and *yjdN* transcription. To explore how *REP325* may affect the operon’s transcription specifically, we took advantage of the simple *yjdMN* operon architecture (two genes flanking the *REP325* element) and used Northern blotting to compare transcript abundance across different operon constructs. We employed ^32^P-labeled DNA probes because radioactive Northern blots provide the sensitivity needed to quantify transcripts over a wide dynamic range and to resolve RNA species with and without the noncoding *REP325* sequence.

We first designed a ^32^P-labeled M-probe specifically targeting *yjdM* transcript. The M-probe detected a robust band at the expected size in *wt* cells overexpressing *yjdM* from plasmid pM (**Fig. 3B**, lane 3). Cells overexpressing the *yjdN* transcript from a plasmid pN (**Fig. 3B**, lane 7) had no detected signal, confirming probe specificity. Endogenous *yjdM* expression from the chromosomal locus was not detected in *wt* cells (**Fig. 3B**, lane 1, **Fig. S11**) or *ΔREP325* cells (**Fig. 3B**, lane 2, **Fig. S11**).

Next, we used the same M-probe to analyze transcripts from plasmids expressing *yjdM-REP325* (pMR), *yjdM-REP325-yjdN* (pMRN), and *yjdM-yjdN* (pMN) under identical conditions. Notably, the predominant product of pMRN did not correspond to the full-length *yjdM-REP325-yjdN* transcript but instead migrated at a size similar to that of pMR (**Fig. 3B**, lane 4 vs. 5), consistent with an RNA containing *yjdM* plus part of *REP325*. A transcript containing *yjdM* and the entire *REP325* would be similar in size to the expected *ydjM-yjdN* transcript and thus longer than the pMR or primary pMRN transcript band (**Fig. S11**). Moreover, the detected transcript bands of pMR and pMRN were broader than that of pM (**Fig. 3B**, lanes 4 and 5 vs. lane 3), suggesting a mixed population of transcripts terminating at multiple positions within *REP325*, likely because the repeated hairpins act as a series of leaky terminators and/or exonuclease RNAse roadblocks.

Strikingly, the *yjdM* transcript level expressed from pMN was more than 160-fold lower than that from pMRN (**Fig. 3B**, lane 5 vs. 6, **Fig. 3B, Table S14**), even though all constructs shared the identical promoter. Assuming similar transcription initiation rates for the *yjdM* portion of pMRN and pMN, this large difference in steady-state *yjdM* mRNA levels between pMRN and pMN is most likely explained by differential degradation of the two transcripts due to the presence and absence of REP325. Additionally, the enhanced *yjdM* mRNA levels in pM, pMR, and pMRN constructs correlated with the prolonged lag phases of these cells.

Finally, *REP325* appeared to enhance *yjdM* accumulation only when *yjdN* was present; we did not observe a significant increase in *yjdM* levels when comparing either pMR or pMRN to pM (**Fig. 3B**, lanes 3 vs. 4 vs. 5, **Fig. 3C, Table S14**). These results suggested that *REP325* protects *yjdM* from degradation primarily in the genetic-context of *yjdN* RNA, which may trigger decay into *yjdM* RNA in the absence of REP325.

### REP325 attenuates transcription upstream of yjdN

Having established that *REP325* stabilizes upstream *yjdM* gene expression in the presence of *yjdN*, we next asked how it affects the downstream *yjdN* gene. Previous work suggested that REPs can function as transcriptional terminators (20), raising the possibility that *REP325* attenuates transcription after *yjdM* and limits *yjdN* transcription. To examine this possibility, we designed a ^32^P-labeled N-probe specifically targeting *yjdN* and probed the same RNA samples used for y*jdM* (M-probe) analysis (**Fig. S11B**, Materials and Methods). The N-probe produced strong, size-appropriate bands for *yjdN* transcripts expressed from pN, confirming the specificity of the probe (**Fig. 3D**, lane 7).

In contrast to our plasmid control, endogenous *yjdN* signal was only detected at low levels in *ΔREP325* cells (see increased contrast images in **Fig. S11C-E**, lanes 1 and 2). Detection of endogenous *yjdN* only in *ΔREP325* cells is consistent with elevated *yjdN* expression in *ΔREP325* as observed by RNA-seq (**Fig. 3A**). The endogenous *ΔREP325 yjdN* transcript length matched that from pMN (*yjdM-yjdN* without *REP325*), confirming that pMN recapitulates *ΔREP325*

When overexpressed from plasmids (**Fig. 3D**, lanes 5 vs.6), *yjdN* transcripts levels from pMN (*yjdM-yjdN* without *REP325*) was ∼ 8-fold higher than that from pMRN (*yjdM–REP325–yjdN*, **Fig. 3E**, **Table S14**), indicating that *REP325* reduced readthrough into *yjdN* and thus acted as an attenuator/terminator between *yjdM* and *yjdN*. Notably, *yjdN* expression increased upon *REP325* removal by the same amount in the Northern blot and RNA-Seq experiments (**Fig. S10 and 3E**).

Combining both sets of Northerns (**Fig. 3B-E**), we found that *REP325* both stabilized upstream *yjdM* mRNA and promoted premature termination before *yjdN*. This dual role, simultaneous stabilization of an upstream transcript and attenuation or termination of a downstream gene, has not, to our knowledge, been previously demonstrated for a REP element.

### REP325 is expressed as a ladder of truncated transcripts

To investigate whether *REP325* alone could indeed function as a transcription attenuator, we designed a ^32^P-labeled probe (R-probe) targeting the second hairpin of the naRNA4 sequence in *REP325* (**Fig. 3F**). This probe detected a strong, specific signal from *in vitro* transcribed (IVT) naRNA4 in 6% denaturing polyacrylamide gels (**Fig. 3F**, lane 6, Material and methods), but no detectable signal in *ΔREP325* cells, confirming probe specificity (**Fig. 3F**, lane 2).

We next used the R-probe to examine *REP325*-derived transcripts from three plasmids: pREP325 (full-length 12-hairpin REP325), pnaRNA34 (four internal hairpins corresponding to naRNA3 and naRNA4), and pnaRNA4 (two hairpins corresponding to naRNA4). To ensure that only the naRNA34 or naRNA4 region was transcribed, we placed an *rrnD* terminator immediately downstream in pnaRNA34 and pnaRNA4 (**Fig. 12A**). All three constructs were robustly transcribed but did not produce any obvious growth phenotype in either *wt* or *ΔREP325* backgrounds (**Fig. S7, Fig. S8**). For pnaRNA4, the shortest band migrated at a similar size as IVT-naRNA4, and a longer doublet matched naRNA followed by the *rrnD* terminator, which has two closely spaced termination sites (**Fig. 3F**, lane 5). For pnaRNA34, we detected a series of bands whose inferred lengths corresponded to transcripts containing ∼ 2, 3, or 4 hairpins, with the longest band matching the four-hairpin naRNA34–*rrnD* product (232 nt, **Fig. 3F**, lane 4). In contrast, pREP325 produced a striking, ladder-like pattern of bands (**Fig. 3F**, lane 3), with highly reproducible spacing across replicates (**Fig. S12B-D**). The most intense band from pREP325 comigrated with the putative four-hairpin pnaRNA34 band and accounted for ∼ 50% of the total signal (**Fig. 3F**, **Fig. S11B-D**, lane 3 arrow). Several additional, progressively longer pREP325 transcripts were detected, but even the longest species was shorter than the expected full-length, 12-hairpin *REP325-rrnD* transcript (**Fig. S11B-D**). Thus, under these conditions, full-length *REP325* RNA was not a major product. Instead, *REP325* is expressed as a ladder of discrete hairpin-containing species generated by transcription termination or mRNA decay.

### REP325 transcription attenuation exhibits partial rho-dependence

Previously, REP*s* have been proposed to act as Rho-dependent terminators (20). Therefore, we asked whether any of these REP325-derived products resulted from Rho-dependent termination. We treated cells with bicyclomycin (BCM), a small-molecule antibiotic that specifically inhibits Rho’s ATPase and translocase activities and therefor blocks Rho-dependent termination. BCM treatment had little effect on the transcript length distributions from pnaRNA4 and pnaRNA34 (**Fig. 3F**, lanes 8 and 9), indicating no detectable Rho-dependence for these shorter REP fragments.

By contrast, pREP325, which lacked a single dominant full-length product, showed a clear response to BCM (**Fig. 3F**, lane 7). Upon Rho-inhibition, we reproducibly observed additional bands at higher apparent molecular weights than the longest species in the untreated sample, as well as a marked increase in the intensity of a band just above the 4-hairpin-like primary product (**Fig. 3F**, lane 7). We quantified transcription read-through for REP325 as the fraction of signal in bands migrating more slowly than the primary four-hairpin-like band (**Fig. S11B-D**). BCM increased read-through from 23±7 % to 39±16%, nearly a two-fold change (**Fig. 3G**). The BCM-induced accumulation of longer transcripts indicated that at least a subset of REP325-mediated termination events were Rho-dependent. However, at the same time, the lack of BCM sensitivity for pnaRNA4 and pnaRNA34 showed that 2–4-hairpin portions of REPs are not always sufficient for strong Rho-dependent termination. In the case of *REP325*, partial Rho-dependence emerges only the full-length *REP325* array.

In support of a role for REPs as Rho-dependent terminators, a previous unbiased *in vitro* screening identified Rho-binding sequences at 78 unique REPs within the *E. coli* genome (22) (**Table S16**). These REPs included 28 REPs that had been previously identified as Rho-dependent 3’ ends *in vivo* but were classified as RNase processing sites based on their C/G ratios (46). The same *in vitro* screen also mapped Rho-binding sites within 100bp of 72 additional unique REPs. However, *REP325* and several other known Rho-binding sites were not recovered in the *in vitro* screen, indicating that efficient Rho engagement at these sites may require additional *in vivo* factors or conditions not recapitulated in the assay (22).

### REP sequence varies at mismatches while REP structure varies in hairpin length

In *E. coli*, all REP elements are extragenic, with ∼ 40% located between tandem genes, ∼ 60% between convergent genes, and ∼ 1% between divergent genes. This strong bias toward tandem and convergent genes suggests that REPs, like *REP325*, may generally act as 3’ UTR-associated regulators of adjacent genes. Because *REP325* dually regulates mRNA decay and transcription termination, we asked how broadly these conclusions might extend to other REPs in the genome.

To explore this possibility, we first quantified sequence and structure conservation for the 697 palindromic units (PUs) that make up all annotated REP elements in the *E. coli* MG1655 reference genome (4). For each PU, we scored how closely it matched the canonical Y, Z1, and Z2 PU hairpin structures and sequence motifs as defined by Ton-Hoang *et al* (3) (**Fig. 4A-B**, color-coded bases, Methods). We found that sequence variations concentrated in the identities of unpaired bases within hairpin bubbles and loops (**Fig. 4A**) whereas structure variations were dominated by differences in the hairpin length (**Fig. 4B**). Furthermore, sequence and structure conservation scores for individual PUs were highly correlated (Tandem Spearman *r* = 0.77, Convergent Spearman *r* = 0.83, **Fig. S13A-B**), suggesting that both features are maintained by similar evolutionary pressures. REPs located between tandem and convergent gene pairs showed similar sequence and structure conservations scores (**Fig. S13C-D**), suggesting that that any distinct effects on tandem vs. convergent genes are not simply due to one class being more or less conserved than the other.

**Figure 4.**
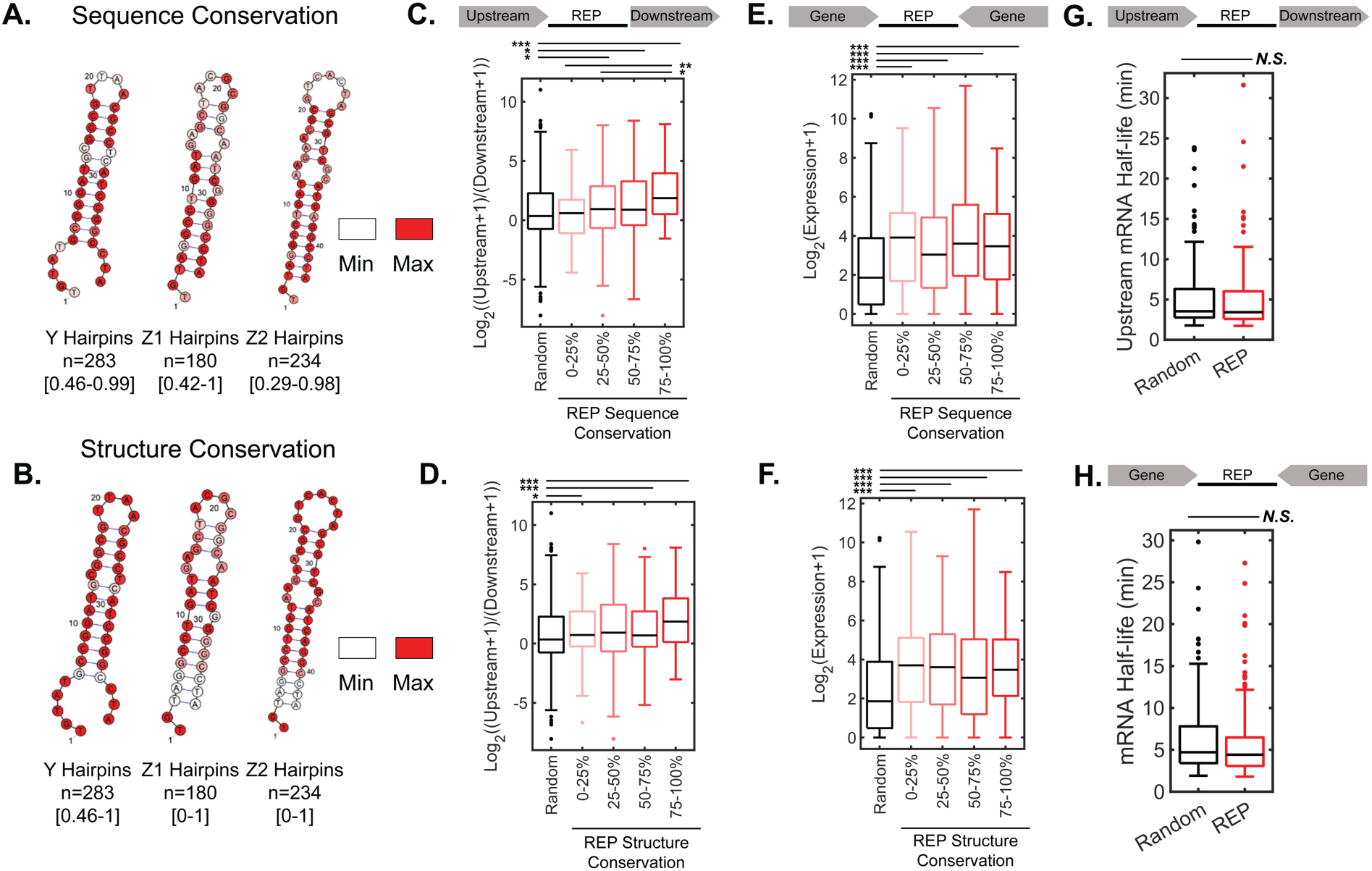
REPs conservation is associated with biased upstream/downstream expression in tandem gene pairs and elevated expression of convergent gene pairs. **A** and **B.** Sequence conservation scores (**A**) and structure conservation scores (**B**) by nucleotide for each PU hairpin consensus sequence. **C and D.** Expression ratio of upstream/downstream genes in the random tandem pair set lacking REPs and in tandem pairs containing REPs split into quartiles by REP sequence conservation (**C**) and structure conservation (**D**). **E and F.** Expression levels of convergent gene pairs in the random pair set lacking REPs and in convergent pairs containing REPs split into quartiles by REP sequence conservation (**E**) and structure conservation (**F**). **G.** mRNA half-lives (29) for upstream genes containing REPs compared to upstream genes lacking REPs (randomly sampled) show no statistically significant differences. **H.** mRNA half-lives (29) for convergent genes containing REPs compared to convergent genes lacking REPs (randomly sampled) showed no statistically significant difference. Significance determined by Wilcox’s rank sum test. * *p*-value<0.05, ** *p*-value<0.01, *** *p*-value<0.001

### REP-containing tandem pairs have elevated upstream/downstream expression ratio

Using the *wt* RNA-seq data collected for earlier experiments, we quantified transcriptional bias between upstream and downstream genes for all tandem gene pairs separated by a REP (146 REP tandem gene pairs) and compared them to those of a random subset of tandem genes lacking a REP (146 of 2451 total pairs). On average, genes upstream of REPs had higher expression than the random upstream genes (two-sided Wilcoxon’s rank sum test *p*-value < 0.05, **Fig. S14A)**. Although genes downstream of REPs did not show lower expression than random downstream genes, the expression of genes downstream of REPs clearly decreased with increasing REP sequence conservation (**Fig. S14B-C**).

Next, we asked whether the upstream to downstream gene expression ratios in REP-containing tandem genes differ from those in the random control sets, and how these ratios vary with REP sequence and structure conservation. When all REP-containing tandem pairs were analyzed together, the upstream/downstream expression ratios were significantly higher than those in random tandem genes (median log-transform ratio = 0.96 in REP-containing genes vs. - 0.09 in random genes, two-sided Wilcoxon’s rank sum test *p*-value < 0.05, **Fig S14D**), consistent with a general tendency of REPs to favor upstream gene expression, as observed for *REP325* in the *yjdMN* operon. However, both REP-containing and random pairs showed a wide spread of ratios, indicating that additional factors likely influence the magnitude of this effect across different genes or locations.

To test whether this expression bias was associated with more canonical REPs, we grouped tandem pairs by the average sequence and structure conservation scores of their intervening REPs and compared the upstream/downstream ratios to those of the random tandem set. We found that upstream/downstream expression ratios increased with REP sequence conservation (Spearman’s correlation ρ = 0.2, *p* = 0.014, **Fig. S14E**), and REPs in the top three sequence conservation quartiles showed significantly elevated ratios compared to the random tandem genes (**Fig. 4C**, two-sided Wilcoxon’s rank sum test *p*-value < 0.05). For structure conservation, first, third, and fourth quartiles of REP-containing genes showed significantly higher upstream/downstream expression ratios than random tandem genes (**Fig. 4D**, two-sided Wilcoxon’s rank sum test *p*-value < 0.05), although the ratio did not correlate significantly with structure conservation (Spearman’s correlation ρ = 0.12, p = 0.15, **Fig. S14F**). Together, these results indicated that biased upstream and downstream expression is a common feature of REP-containing tandem pairs, particularly for REPs with highly conserved sequences and sufficient structure conservations.

### REP-containing convergent genes show elevated transcription

Between convergent genes (**Fig 4E**), a palindromic REP could function bidirectionally as both a terminator and a degradation block, thereby increasing the steady-state abundance of both transcripts. In *E. coli*, ∼ 30% of convergent gene intergenic regions contain a REP (207/704 gene pairs). Using the same RNA-seq dataset of *wt* cells, we compared transcription levels for all convergent gene pairs with an intervening REP to a random subset of convergent pairs lacking a REP (**Fig. S14G**). We observed that convergent gene pairs associated with REPs showed significantly higher expression than random pairs across all quartiles of REP-containing genes stratified by REP sequence or structure conservations (**Fig. 4E-F**), but expression did not correlate significantly with either conservation scores (**Fig. S14H-I**). Thus, even moderately canonical REPs can enhance expression of convergent genes; this effect appears to depend primarily on REP presence rather than fine differences in conservation.

To further examine the idea that REPs act as bidirectional terminators between convergent genes, we took advantage of a previous sequence-based study that identified 306 bidirectional terminators between convergent genes in the *E. coli* genome (47). We found 78 REPs within these terminators, accounting for nearly 40% of all REPs between convergent genes (**Table S16**).

### Terminal REPs are not associated with longer endogenous mRNAs half-lives

Our results so far showed that *REPs* favored upstream gene transcription in tandem pairs and enhanced expression of both genes in convergent pairs. Based on our Northern blot results, we asked whether these effects could be explained by increased mRNA stability. If so, mRNAs from genes with terminal REPs might be expected to have longer half-lives than those from genes lacking REPs. To test this possibility, we analyzed a published dataset that measured mRNA half-lives in rifampicin-treated *E. coli* cells (29). However, contrary to this expectation, genes with a terminal REP showed no significant difference in half-lives than a random set of genes with similar expression levels lacking REPs (**Fig. 4G**). We also observed no significant difference in half-lives for convergent genes with REPs compared to expression-matched random control (**Fig. 4H**). These observations suggest that terminal REPs are not globally associated with increased transcript stability at endogenous expression levels.

## Discussion

In this work, we demonstrated that REPs in *E. coli* do not make a major contribution to long-distance chromosome organization but instead act as local, 3-UTR-associated transcription regulators of neighboring genes. Our data supports a model in which REPs modulate both transcription termination and mRNA decay in a context-dependent manner, providing a unifying framework that reconciles previously varying observations about REP-dependent gene regulation. While our results do not preclude additional roles for REPs in processes such as supercoiling control (8, 48) or translation regulation (9), our genome-wide analysis of REP-adjacent genes indicates that their predominant impact *in vivo* is to fine-tune local transcription across the genome.

### HU-naRNA4 does not compact the chromosome

Our results do not support a major role for HU-naRNA4 in chromosome compaction as previously proposed (12–14). We observed no appreciable HU-naRNA-crfDNA complex formation (**Fig. 1**), no changes in nucleoid morphology upon deleting *REP325* or overexpressing REP RNAs, and no clear pattern of REP-REP contacts in available Hi-C datasets (**Fig. 2**). Although differences in cell growth condition (minimal medium in prior studies vs. rich defined medium here) could contribute to the discrepancy, our results are also consistent with very low endogenous transcription of *REP325* RNAs (on the order of ∼1-2 copies per cell, Supplementary Information, **Fig. S11B-D**). In addition, our previous single-molecule tracking showed that *REP325* deletion did not alter HU’s DNA binding dynamics in live cells (25), and independent studies have reported minimal effects of *REP325 (nc5)* deletion on chromosome structure (49, 50). Taken together, these observations suggested that the proposed HU–naRNA4–interactions are, at best, not a robust mechanism of chromosomal compaction.

### Unresolved functions of yjdM and yjdN

The growth-rate phenotypes we observed in the *ΔREP325* and *yjdM* overexpression strains are intriguing but also puzzling, because our whole-cell RNA-seq analyses did not reveal a clear, pathway-level explanation (**Fig. S10**). The simplest interpretation is that these phenotypes arise from the specific functions of *yjdM* (and possibly *yjdN*), yet the molecular roles of both genes remain poorly defined. Existing literature on *yjdM* and *yjdN* is conflicting (39, 40), and a recent CRISPRi screen reported no fitness effect for either gene across fifteen *E. coli* strains (51). Our results instead indicated that altered *yjdM* expression affect cell viability in the stationary phase and/or lag-phase exit (**Fig. S9**). Thus, the absence of a clear theme among *yjdM*-associated differentially expressed genes is perhaps not surprising, given that our RNA-Seq measurements were collected from exponential phase cells. A direct comparison of stationary-phase transcription profiles between *wt* and *yjdM*-overexpression cells would therefore be an interesting direction for future work.

### REP325 acts as a dual terminator–stabilizer in the yjdMN operon in a context-dependent manner

Our RNA-Seq data (**Fig 3A)** and Northern blot results (**Fig. 3B-E**) together showed that REP325 plays a dual regulatory role in the y*jdMN* operon, simultaneously promoting *yjdM* stability and limiting *yjdN* transcription. Under both overexpression and endogenous conditions, removing *REP325* increased the downstream *yjdN* expression by approximately one order of magnitude (Northern blot: ∼8-fold for pMRN vs. pMN **Fig. 3D**; RNA-Seq: ∼10-fold for *wt* vs. *ΔREP325* **Fig. S12A**), consistent with a robust termination/attenuation function that is largely independent of the expression level. In contrast, the impact of *REP325* on the upstream yjdM was strongly expression-dependent: *REP325* removal caused ∼160-fold decrease in *yjdM* signal under the overexpression condition (pMRN vs. pMN, **Fig. 3A**), but only ∼3-fold at the endogenous locus as measured by RNA-seq (*wt* vs. *ΔREP325*, **Fig. S12A**). These results imply that REP-mediated stabilization effect could be markedly amplified at higher transcriptional output. Although some variation is expected between Northern blot and RNA-seq measurements, the close agreement for *yjdN* (∼8–10-fold increase in both assays) argues that the >100-fold difference in apparent *yjdM* stabilization cannot be explained by methodology alone.

One interesting observation is that *REP325* specifically stabilized *yjdM* in the presence of *yjdN* but not when *yjdM* was expressed alone. In plasmid constructs, including *REP325* (pMR or pMRN) did not further increase *yjdM* transcription levels relative to pM, yet all three had ∼160-fold more *yjdM* mRNA than the pMN construct, even though pM and pMN shared the same promoter and 3′ UTR and neither contained *REP325* sequence. This large difference between pM and pMN mRNA levels suggests that *yjdM* sequence alone encodes an intrinsically stable transcript and/or that the downstream *yjdN* transcript is unusually unstable, despite lacking obvious sequence features that might trigger quality-control pathways. Such context-specific stabilization by could explain why we do not see a global increase in mRNA half-life at *REP*-containing termini. It also remains possible that *REPs* preferentially insert downstream of intrinsically unstable transcripts; rigorously testing this hypothesis will require direct half-life measurements for each gene with and without its associated REP.

### Genome-wide RNA-seq supports REPs’ roles as terminators and stabilizers

Our study is, to our knowledge, the first to systematically relate intragenomic sequence and structure conservations of REPs to neighboring gene expression using RNA-seq. We found that REPs with more canonical sequence and Y/Z-type hairpin structures were more frequently associated with higher upstream/downstream expression ratios in tandem gene pairs, consistent with REP-mediated termination and protection of upstream transcripts. Additionally, convergent genes with intervening REPs were expressed at higher levels than convergent pairs lacking REPs, consistent with REPs acting as bidirectional terminators and/or degradation blocks that stabilize transcripts on both sides of the intergenic region. At the same time, the broad spread in the expression ratios indicates that other factors, such as local expression level, intrinsic transcript stability, and surrounding sequence context, may substantially modulate the magnitude of REP regulatory effects.

REPs are rarely highlighted in *in vivo* genome-wide screens of transcription termination (21, 47) or mRNA decay (52), but this omission likely reflects REPs’ dual roles rather than a lack of their involvement in the two processes. For example, a study of mRNA decay regulation explicitly removed sites implicated in Rho-dependent termination (52), whereas termination-focused studies commonly discarded 3’ ends affected by deletion of RNA-degradation factors (21), including factors known to process REP-mediated degradation (18, 53). Specifically, Peters *el al.* retrospectively labeled many REPs as mRNA decay sites based on their roughly equal C:G content, even though about half of REPs lie within 300bp of a Rho-dependent 3’ end (21). When we queried REPs explicitly in a Rho-binding screen and termination study (22, 47), we did find overlaps with REPs. Library- or SELEX-based *in vitro* screens can offer a cleaner approach to map which termination or degradation factors bind to REPs, but targeted *in vivo* studies will ultimately be required to define the specific molecular interactions by which REPs affect transcription and mRNA degradation.

### Possible origins and mobility of REP elements

The variability in REP effects likely reflects how and where they are inserted. REPs have been proposed to be “domesticated transposons,” derived from insertion sequences (IS) that have been evolutionarily co-opted by the host for beneficial functions (3). Although the *in vivo* mechanism of REP insertion is not fully established, the REP-associated tyrosine transposase RAYT cleaves REP hairpins in a structure-specific manner reminiscent of TnpA enzymes on IS elements (3, 54, 55), suggesting a transposition-like mechanism for REP insertion. Semi-random insertion of REP into accessible DNA regions would make REP effects highly context dependent, with local sequences and chromosomal environment influencing termination efficiency and/or degradation blocking. Transposition-like insertion could also explain REP *ori*-proximal overrepresentation and *ter* underrepresentation. A bacteria-adapted ATAC-seq (56) approach would help test this correlation directly. At the same time, REPs themselves are not autonomous mobile elements: *REP325*, *naRNA34*, and *naRNA4* were all stable on plasmids in our experiments, and the high sequence homology of REPs generally argues for active conservation and selection after insertion by some unknown mechanism (7, 57), with REPs retained at locations where their local expression effects are beneficial and lost where they are deleterious.

### REPs as tunable, noncoding dials on gene expression

Our findings suggest REPs can serve as modular, noncoding elements for tuning gene expression without altering coding sequences or core promoters. The MG1655 *REP* landscape represents just one instance of many possible REP distributions; across strains and species, gain or loss of REPs at different 3’ UTR sites could alter expressed protein copy numbers while preserving amino acid sequence, providing a flexible handle for regulatory diversification. Because REP insertion or removal alters termination and mRNA stability in a context-dependent manner, controlled manipulation of REP positions and architectures could be explored to further elucidate their functions and to fine-tune gene expression over a wide range, thereby probing the evolutionarily-tolerated bounds on protein copy number. Based on our data, we expect that REP insertion between genes whose expression is tightly constrained is likely to be deleterious and selected against. However, bacteria can tolerate surprisingly large expression changes even in some essential genes, suggesting substantial room for REP-mediated tuning (58). These considerations raise the exciting possibility of using engineered REP insertions or rearrangements as a noncoding “dial” to adjust expression in synthetic and evolutionary experiments (59), offering a powerful way to interrogate both REP biology and the robustness of bacterial gene regulatory networks.

In conclusion, our work shows that REPs function primarily as local 3′ UTR regulators rather than global chromosomal organizers, modulating neighboring gene expression by jointly influencing mRNA degradation and transcription termination in a context-dependent manner.

## Supporting Information

Supplementary Figures 1-14, Supplementary Information, Tables 1,3,5-7,10, and 11 are attached as Supplementary Information.

Tables 2,4,8,9,12-15 are attached as Supplementary Data.

## Data Accessibility

Relevant code is uploaded on Zenodo. DOI 10.5281/zenodo.19150984

RNA-Seq fastq files have been uploaded to GEO under GSE325648.

## Acknowledgements

This work was funded by NIH 2R35GM136436 to J. X. We thank all Dr. Sarah Woodson’s lab members, including Dr. Yi-Lan Chen and Dr. Roumita Moulick for their advice and training on Northern blots. We also thank LaToya Roker from the Johns Hopkins School of Medicine Microscope Facility for assistance and usage of the OMX-SR structured illumination microscope. This work was carried out at the Advanced Research Computing at Hopkins (ARCH) core facility (rockfish.jhu.edu), which is supported by the National Science Foundation (NSF) grant number OAC1920103. AI Language Models were used to generate initial code used for most data analysis. AI-generated code was thoroughly checked and edited for accuracy by the authors.

