## Supplementary Information for "Repetitive extragenic palindrome (REP) elements are local, context-dependent, dual 3’UTR regulators in *Escherichia coli*"

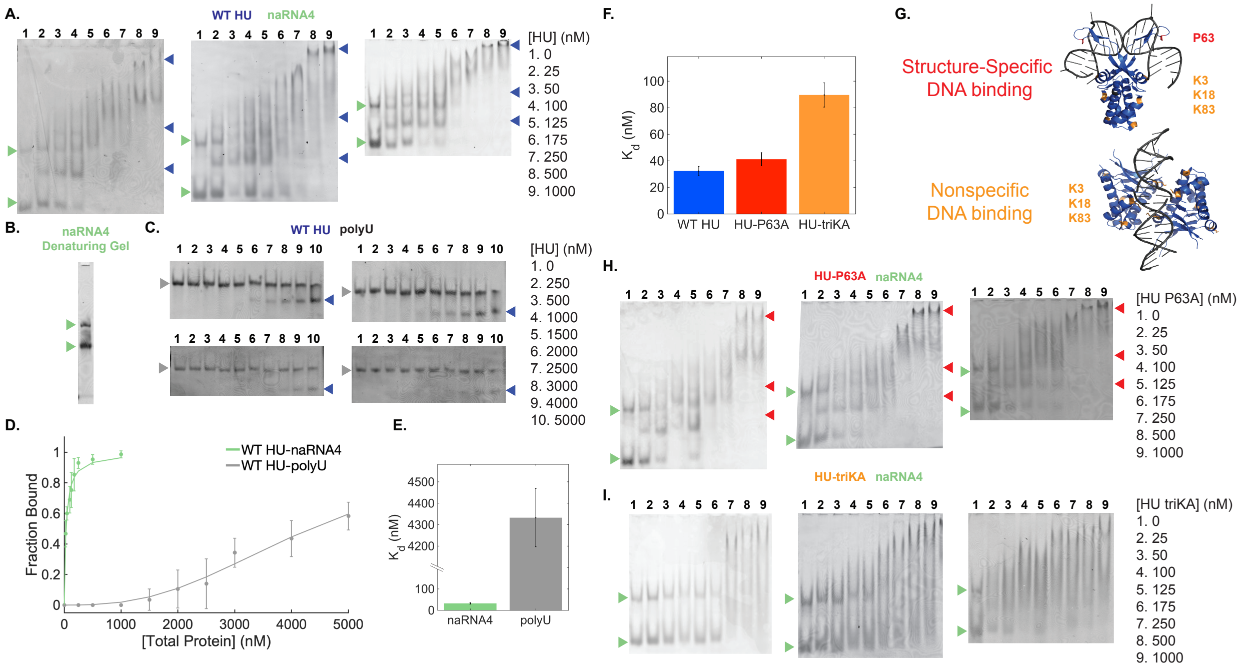

**Supplementary Figure 1. Replicates of HU-RNA binding EMSAs.** All EMSAs are 0.5xTBE 6% polyacrylamide gels with RNA visualized by SYBR™ Gold. **A.** WT HU-naRNA4 EMSAs (n=3 replicates). **B.** 0.5xTBE 6% polyacrylamide 8M Urea Denaturing gel with naRNA4. **C.** WT HU-polyU EMSAs in quadruplicate. **D.** EMSA Quantifications for HU WT binding to naRNA4 (n=3 replicates, green – same data as blue curve in Fig 1C) and to polyU (n=4 replicates, gray). **E.** Quantifications of EMSAs for HU WT binding to naRNA4 and polyU. Data was fit to the Hill equation. Error bars are standard deviation from the mean. **E.** $K_d$ of Hill Equation fits in D. Error bars are standard errors of the fits. **F.** $K_d$ of Hill Equation fits in Fig 1C, error bars are standard errors of the fits (blue bar is same data as the green bar in E.) **G.** Schematics of two HU binding modes (PDB: 4QJU, 4YFT respectively) with conserved proline (red) and important surface lysines (orange) highlighted. **H.** HU P63A-naRNA4 EMSAs in triplicate (n=3 replicates). **I.** HU triKA-naRNA4 EMSAs (n=3 replicates).

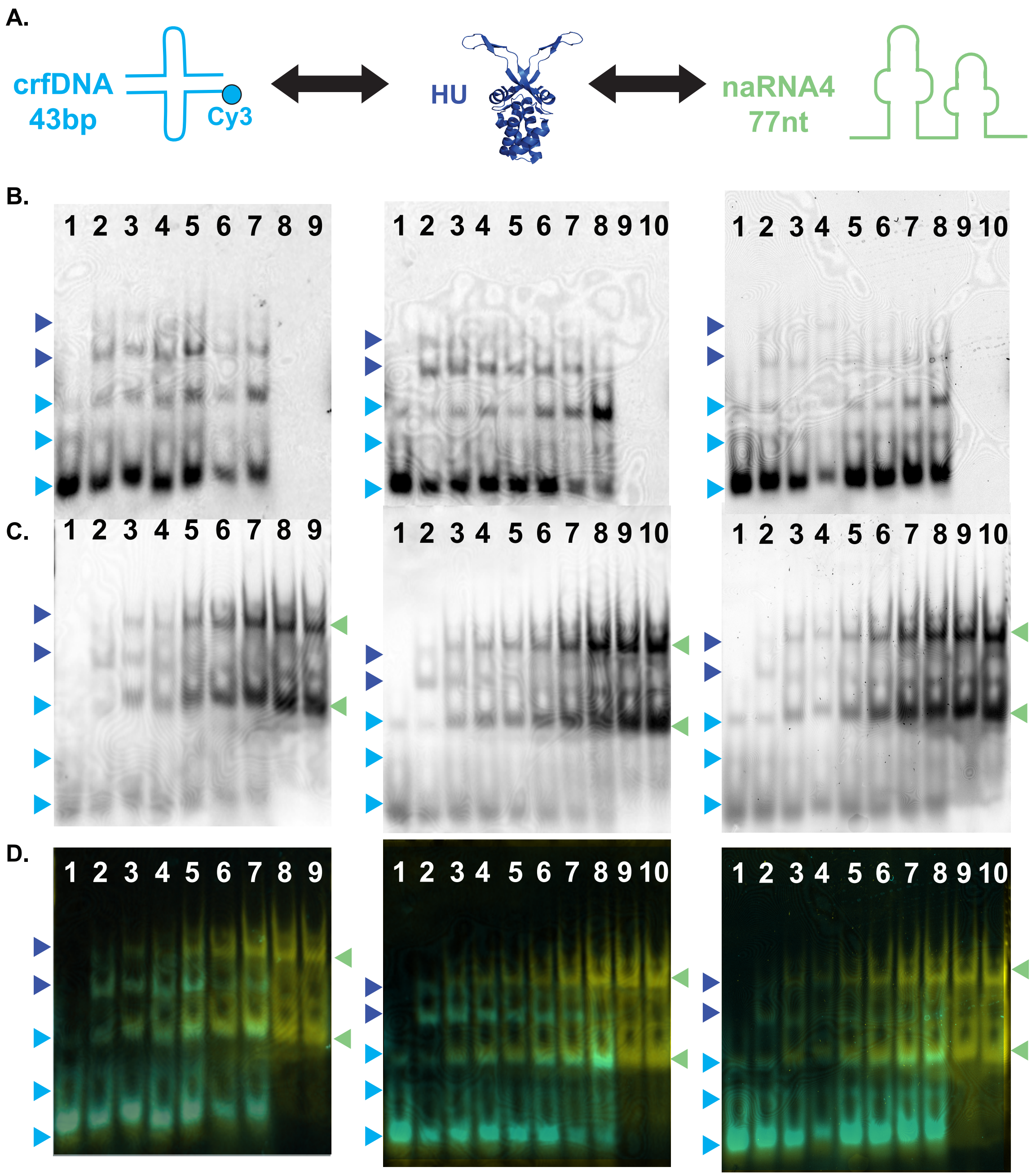

Cy3
SYBR™ Gold

Lanes
1. crfDNA
2. crfDNA: HU
3. crfDNA: HU: 6.25nM naRNA4
4. crfDNA: HU: 12.5nM naRNA4
5. crfDNA: HU: 25nM naRNA4
6. crfDNA: HU: 50nM naRNA4
7. crfDNA: HU: 100nM naRNA4
8. HU: 100nM naRNA4
9. 100nM naRNA4
[crfDNA] = 25nM
[HU] = 50nM

Lanes
1. crfDNA
2. crfDNA: HU
3. crfDNA: HU: 6.25nM naRNA4
4. crfDNA: HU: 12.5nM naRNA4
5. crfDNA: HU: 25nM naRNA4
6. crfDNA: HU: 50nM naRNA4
7. crfDNA: HU: 100nM naRNA4
8. crfDNA: HU: 200nM naRNA4
9. HU: 200nM naRNA4
10. 200nM naRNA4
[crfDNA] = 25nM
[HU] = 50nM

Lanes
1. crfDNA
2. crfDNA: HU
3. crfDNA: HU: 6.25nM naRNA4
4. crfDNA: HU: 12.5nM naRNA4
5. crfDNA: HU: 25nM naRNA4
6. crfDNA: HU: 50nM naRNA4
7. crfDNA: HU: 100nM naRNA4
8. crfDNA: HU: 200nM naRNA4
9. HU: 200nM naRNA4
10. 200nM naRNA4
[crfDNA] = 25nM
[HU] = 50nM

**Supplementary Figure 2. Ternary EMSAs do not show unique crfDNA complex in presence of constant HU and increasing naRNA4. A.** Schematics of crfDNA labeled with Cy3 (cyan) and naRNA4 (green) both of which can bind HU (blue) independently. **B.** 5' Cy3-labeled crfDNA in competition EMSA. Unbound crfDNA (cyan arrows) and HU-bound crfDNA (blue arrows) are indicated. **C.** SYBR™ Gold labeling of naRNA4 and crfDNA in competition EMSA. Unbound naRNA4 (green arrow) is indicated. HU-bound naRNA4 is not observed at these concentrations. **D.** Merged images of the Cy3 (cyan) and SYBR™ Gold (yellow) labeling.

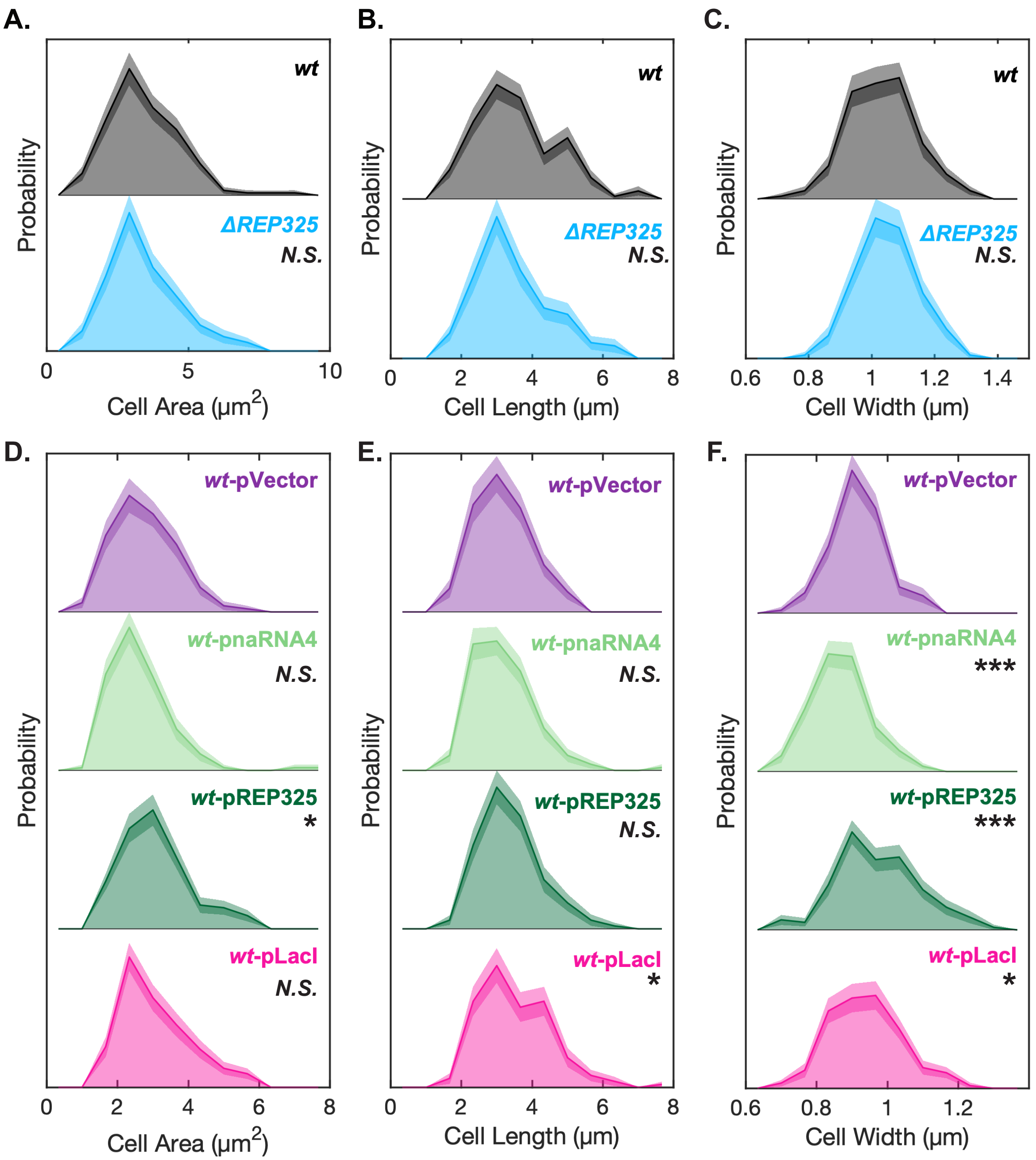

**Supplemental Figure 3. Cell dimensions for conditions in Figure 2. A.-C.** Cell area, cell length, and cell width distributions for *wt* and *ΔREP325* respectively. The two conditions do not significantly differ in cell dimensions. **D.-F.** Cell area, cell length, and cell width distributions for *wt*-pVector (1mM IPTG), *wt*-pnaRNA4 (1mM IPTG), *wt*-pREP325 (1mM IPTG), and *wt*-pLacI (0mM IPTG) conditions. Differences in cell area appear to be primarily explained by differences in cell width. All significant differences between conditions and *wt*-pVector determined by two-sided KS-test. * *p*-value<0.05, ** *p*-value<0.01, *** *p*-value<0.001.

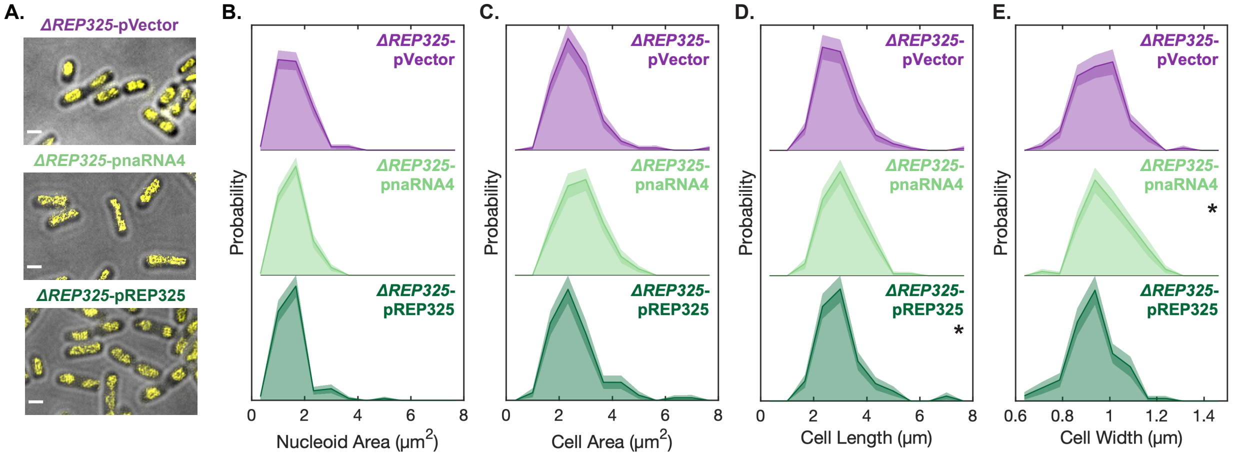

**Supplemental Figure 4. Exogenous expression of REP325 or naRNA4 has no effect on ΔREP325 nucleoid morphology.** *ΔREP325* expressing pVector, pnaRNA4, and pREP325 (all 1mM IPTG) were imaged and quantitatively compared. **A.** Representative SIM images of *wt* and *ΔREP325* cells (gray) with nucleoids stained with the Hoechst dye (yellow). Scale bar: 1 μm. **B-E.** Nucleoid area, cell area, cell length, and cell width distributions, respectively. Only marginally significant differences in cell length and cell width were detected. All other comparisons to *ΔREP325*-pVector were not statistically significant. All significant differences between conditions and *ΔREP325*-pVector determined by two-sided KS-test. * *p*-value<0.05

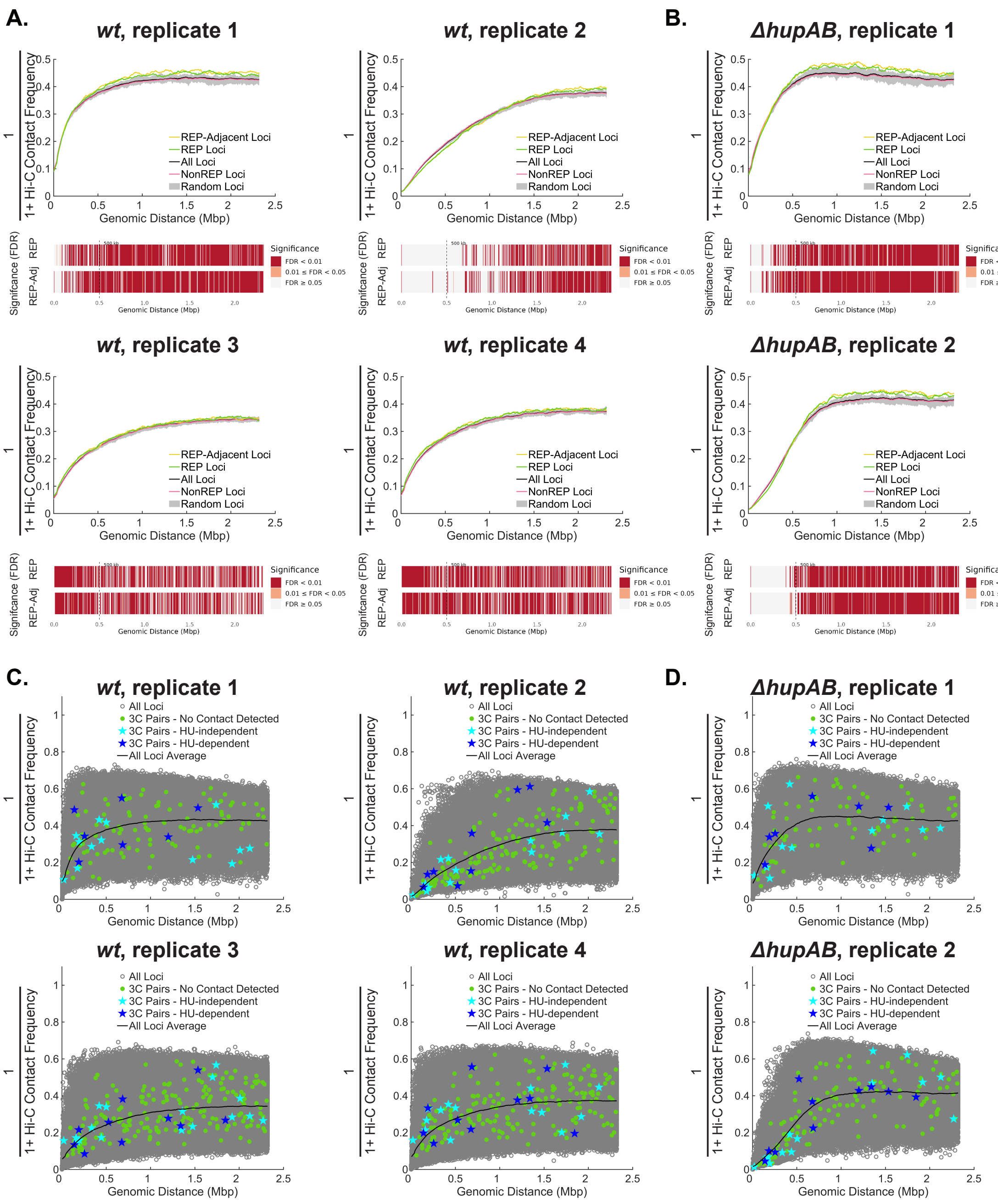

**Supplemental Figure 5. Hi-C data does not show long-distance REP-REP contacts.** Data analyzed from (1, 2). **A.** Hi-C analysis for *wt* condition (N=4 replicates). Average inverse contact frequency vs genomic distance, smoothed with a sliding window average across 15 bins (bin size, 5kb), with same subsets of bins as in Figure 2D. FDR for REP and REP-adjacent loci compared to Random loci plotted beneath. **B.** Hi-C analysis for *ΔhupAB* condition (N=2 replicates). Average inverse contact frequency vs genomic distance, smoothed with a 15-point window. FDR for REP and REP-adjacent loci compared to Random loci plotted beneath. **C.** Specific contact frequencies for REP pairs tested and categorized in Qian *et al* 2015 (3) compared to the contact frequencies of all genomic pairs. **D.** Specific contact frequencies for REP pairs tested and categorized in Qian *et al* 2015 (3) compared to the contact frequencies of all genomic pairs in the *ΔhupAB* conditions.

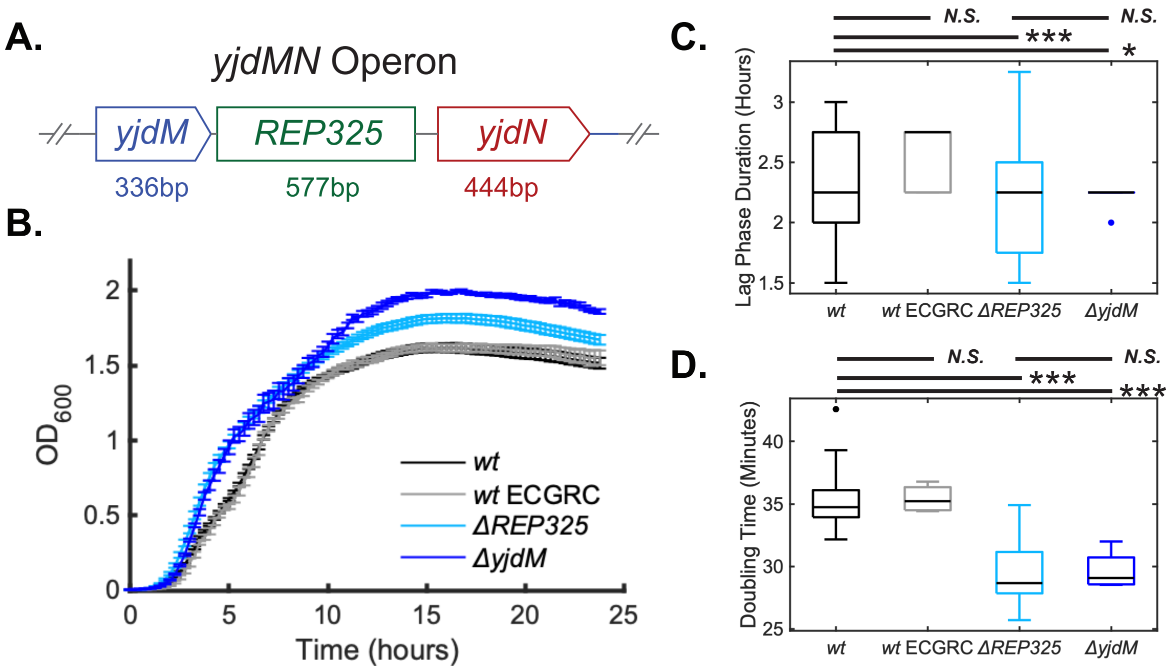

**Supplemental Figure 6. Deletion of *REP325* affects expression of neighboring genes. A.** Schematic of *yjdMN* Operon that contains REP325. Spacing and element lengths are drawn to scale. **B.** Growth curves of *wt*, *wt*-ECGRC*, ΔREP325*, and *yjdM* KO strains (N= 30, 3, 37, and 3 biological replicates respectively, μ±S.E.M.) **C.** Lag phase duration (time from dilution to $OD_{600}$ 0.05) for *wt*, *wt*-ECGRC*, ΔREP325*, and *yjdM* KO **D.** Doubling time (exponential fits of 2.5 hours following $OD_{600}$ 0.05) for *wt*, *wt*-ECGRC*, ΔREP325*, and *yjdM* KO. Growth parameter significant differences determined by two-sided t-test. * *p*-value<0.05, ** *p*-value<0.01, *** *p*-value<0.001

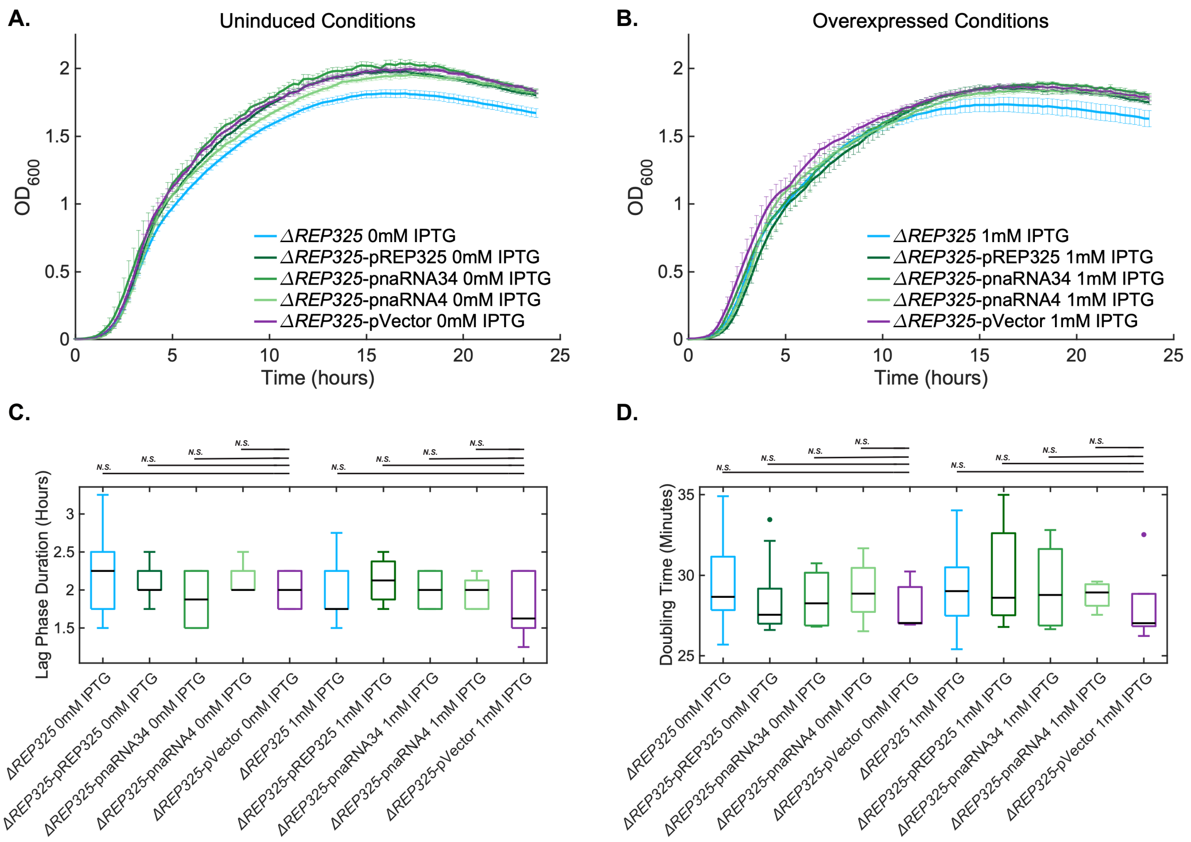

**Supplemental Figure 7. REP expression does not affect growth in *ΔREP325* background A.** Growth curves of *ΔREP325*, *ΔREP325*-pREP325, *ΔREP325*-pnaRNA34, *ΔREP325*-pnaRNA4, and *ΔREP325*-pVector with 0mM IPTG (N= 37, 5, 2, 4, and 3 biological replicates respectively, μ±S.E.M.) **B.** Growth curves of *ΔREP325*, *ΔREP325*-pREP325, *ΔREP325*-pnaRNA34, *ΔREP325*-pnaRNA4, and *ΔREP325*-pVector with 1mM IPTG (N= 18, 4, 2, 4, and 3 biological replicates respectively, μ±S.E.M.) **C.** Lag phase duration for conditions in A. and B. **D.** Doubling time for conditions in A. and B. No significant differences were detected. Growth parameter significant differences *p*-value<0.05, ** *p*-value<0.01, *** *p*-value<0.001

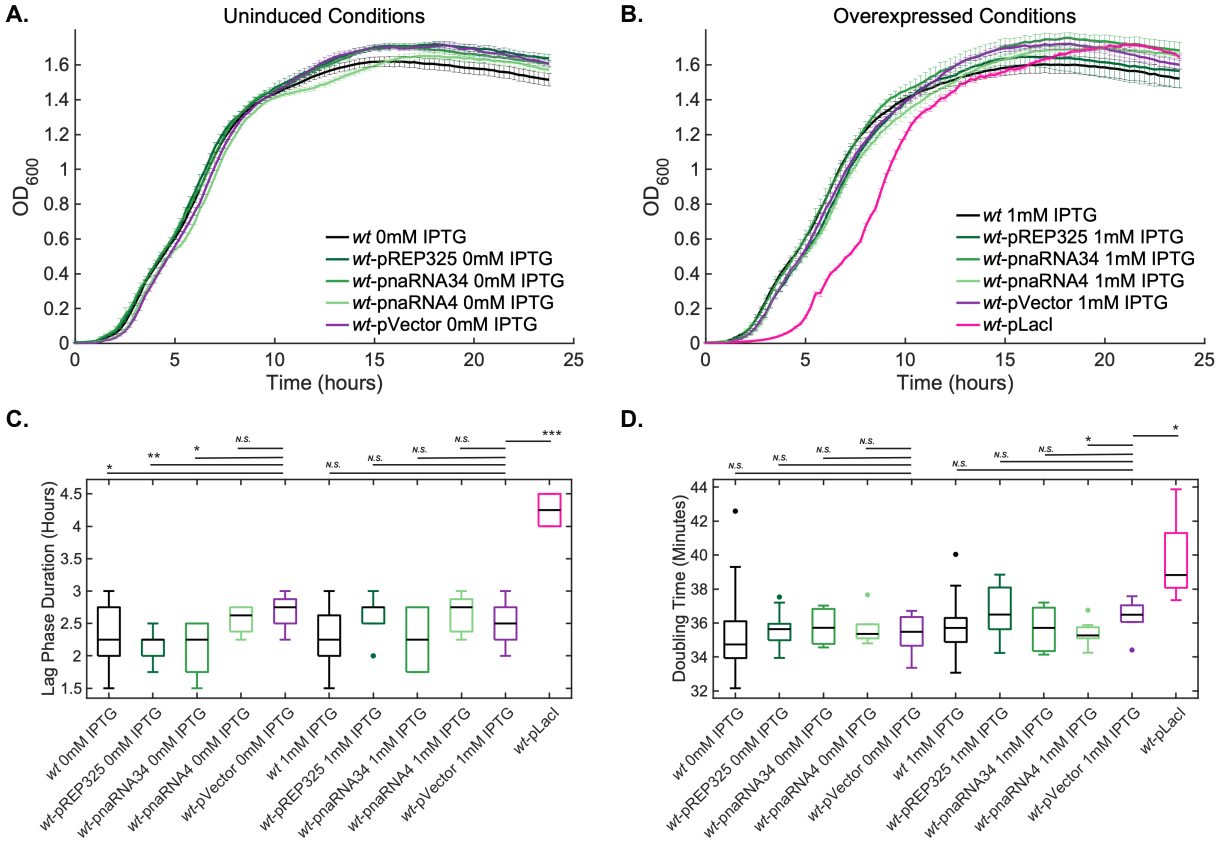

**Supplemental Figure 8. REP expression does not affect growth in *wt* background. A.** Growth curves of *wt*, *wt*-pREP325, *wt*-pnaRNA34, *wt*-pnaRNA4, and *wt*-pVector with 0mM IPTG (N= 30, 5, 3, 4, and 4 biological replicates respectively, μ±S.E.M.) **B.** Growth curves of *wt*, *wt*-pREP325, *wt*-pnaRNA34, *wt*-pnaRNA4, and *wt*-pVector with 1mM IPTG and *wt*-pLacI with 0mM IPTG (N= 18, 5, 3, 4, 4, and 3 biological replicates respectively, μ±S.E.M.) **C.** Lag phase duration for A. and B. **D.** Doubling time for A. and B. Growth parameter significant differences determined by two-sided t-test. Only LacI overexpression significantly slowed growth. * $p$-value<0.05, ** $p$-value<0.01, *** $p$-value<0.001

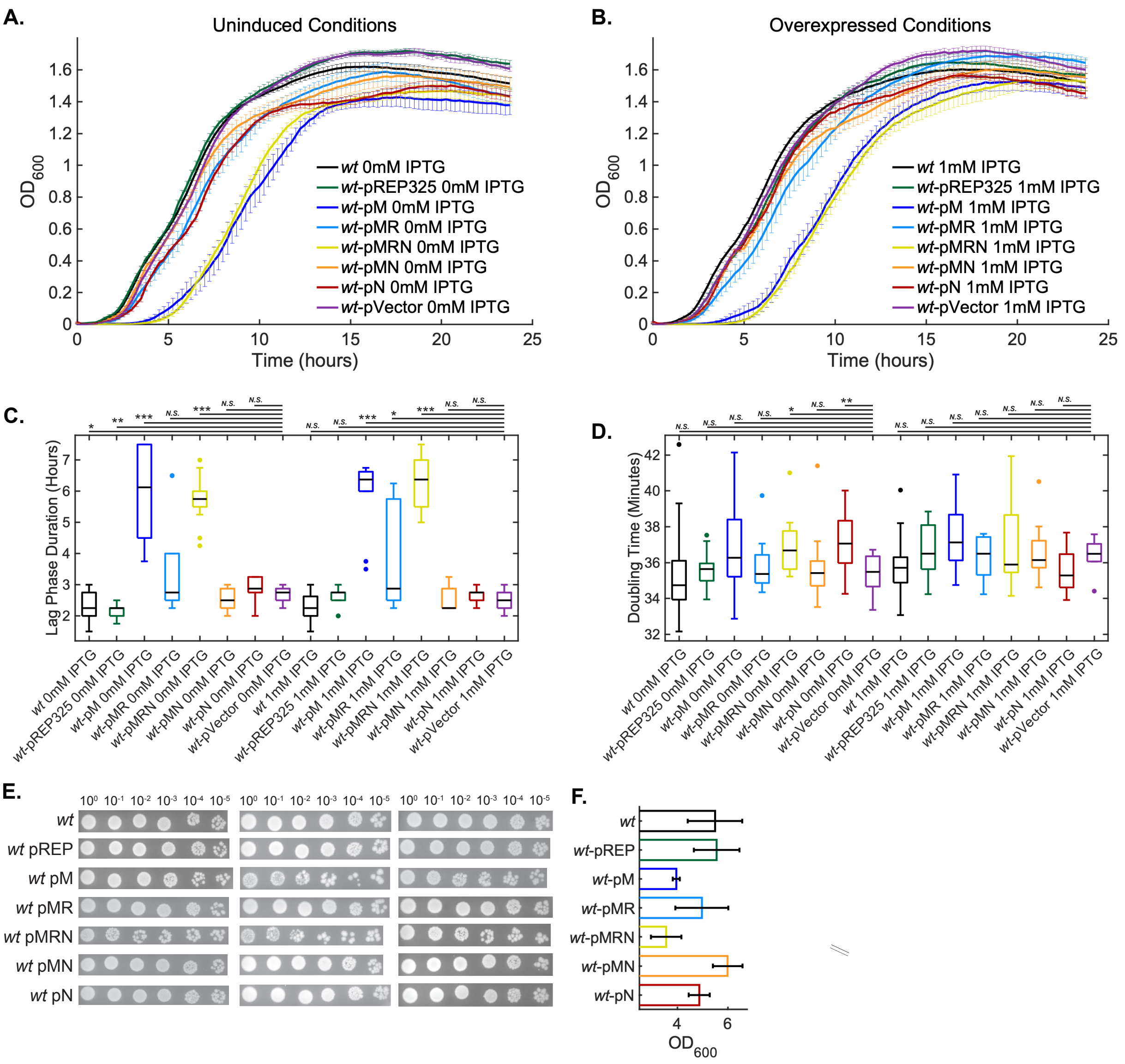

**Supplemental Figure 9. Expression of some *yjdM*-containing mRNAs is associated with extended lag phase and lower stationary phase viability. A.** Growth curves of *yjdMN* operon conditions with 0mM IPTG: *wt*, *wt*-pREP325, *wt*-pM, *wt*-pMR, and *wt*-pMRN, *wt*-pMN, *wt*-pN, and *wt*-pVector (N= 37, 5, 6, 7, 7, 6, 6, and 3 biological replicates respectively, μ±S.E.M.) **B.** Growth curves of *yjdMN* operon conditions with 1mM IPTG (N= 18, 4, 6, 7, 7, 6, 6, and 3 biological replicates respectively, μ±S.E.M.) **C.** Lag phase duration for A. and B. **D.** Doubling time for A. and B. Growth parameter significant differences determined by two-sided t-test* *p*-value<0.05, ** *p*-value<0.01, *** *p*-value<0.001 **E.** Spot dilutions of saturated cultures for *yjdMN* operon conditions without induction (pREP325 abbreviated as pREP). **F.** Concentrations ($OD_{600}$) for saturated cultures used in spot dilutions in E.

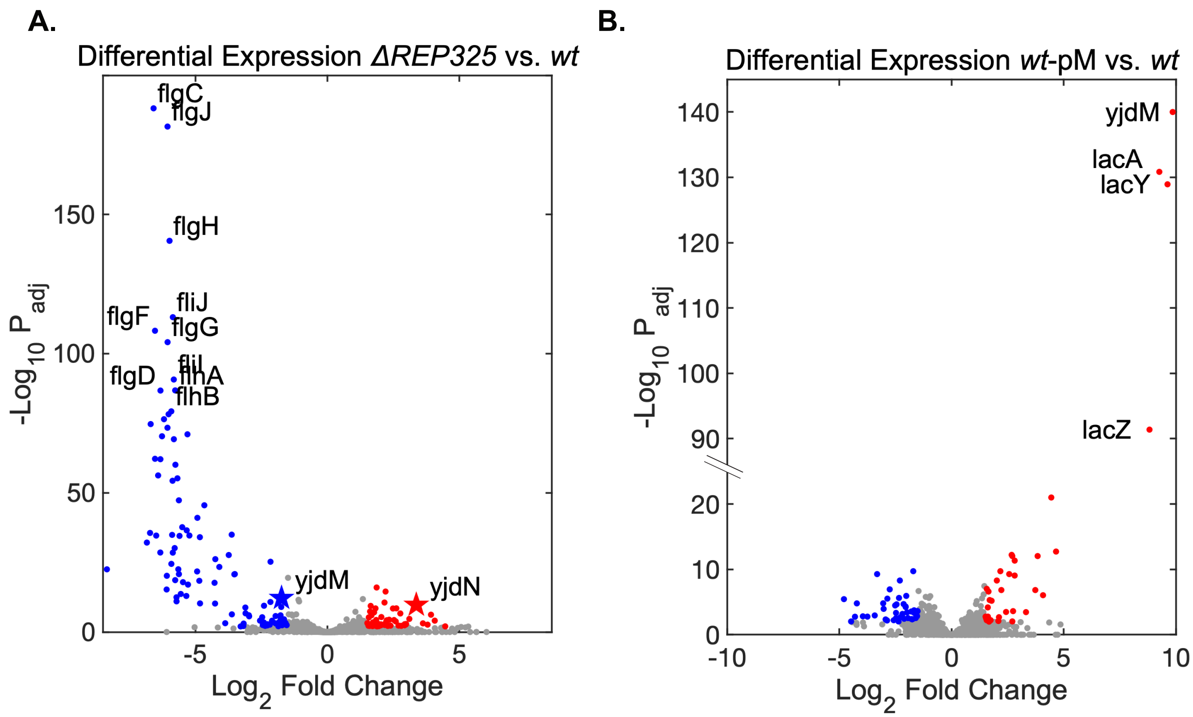

**Supplemental Figure 10. RNA-Seq differential expression analysis** Volcano plots of differential expression analyses. Significantly upregulated genes (red) and significantly downregulated genes (blue) are indicated ($|Log_2FC|>1.5$ and $p_{adj}<0.01$). N=3 biological replicates. **A.** *ΔREP325* vs *wt.* REP-adjacent genes marked with star. **B.** *wt*-pM vs *wt*.

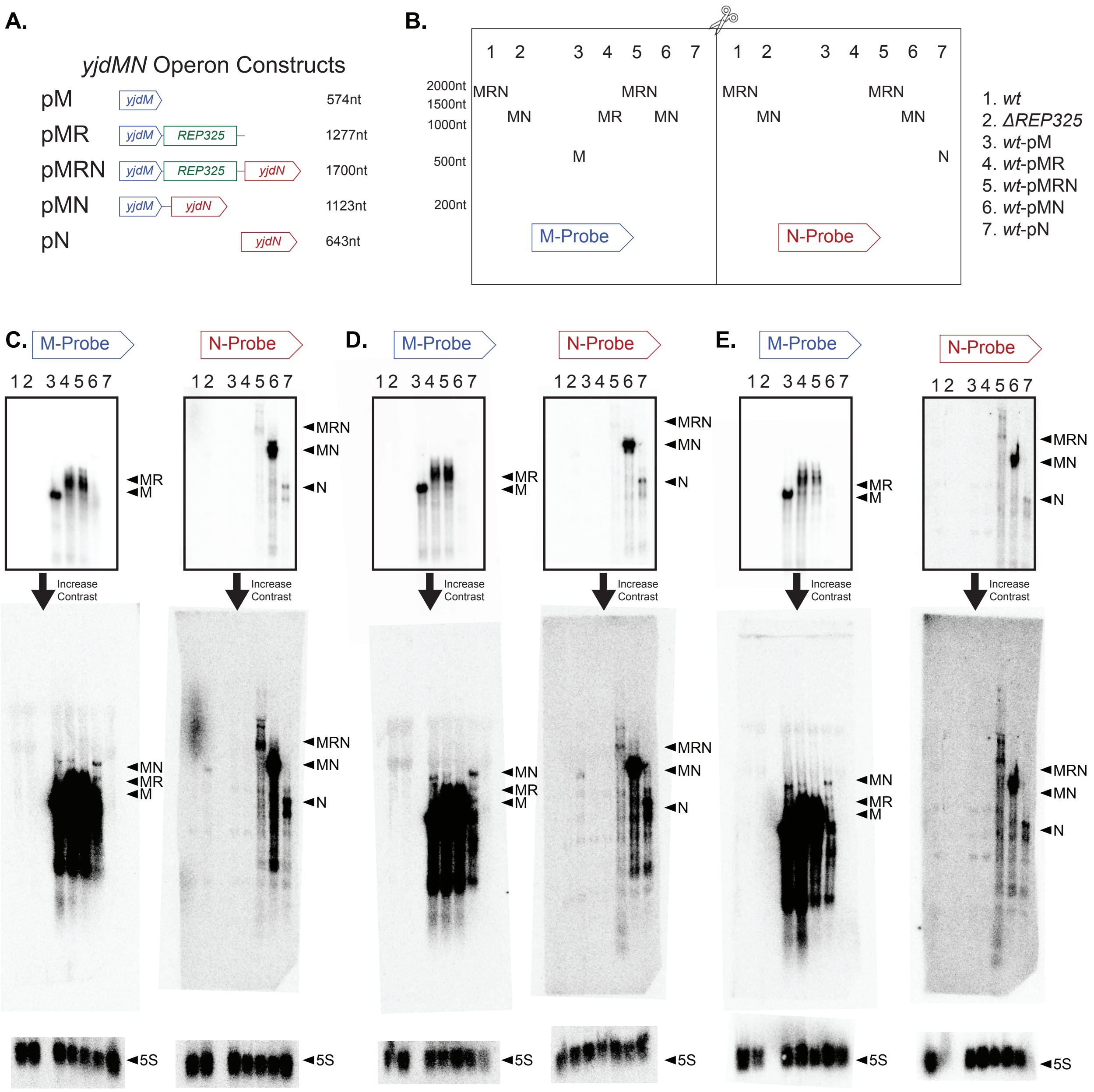

**Supplemental Figure 11. Replicates of *yjdMN* operon Northern blots. A.** Schematic of *yjdMN* Operon Plasmids with maximum transcript lengths **B.** Schematic of Northern Blot with expected full-length bands indicated. **C.-E.** Biological replicates of *yjdMN* operon Northern blots. The same RNA preps were run in duplicate on a single gel and transferred to a single membrane that was then cut in half and probed for *yjdM* and *yjdN* respectively. The *yjdM* mRNA in *wt*-pMN, *yjdN* mRNA in *ΔREP325*, and *yjdN* mRNA in *wt*-pMRN bands are most clearly visualized in the higher contrast images (second row). 5µg (A.), 5µg (B.), and 3µg (C.) of total RNA were loaded for each lane.

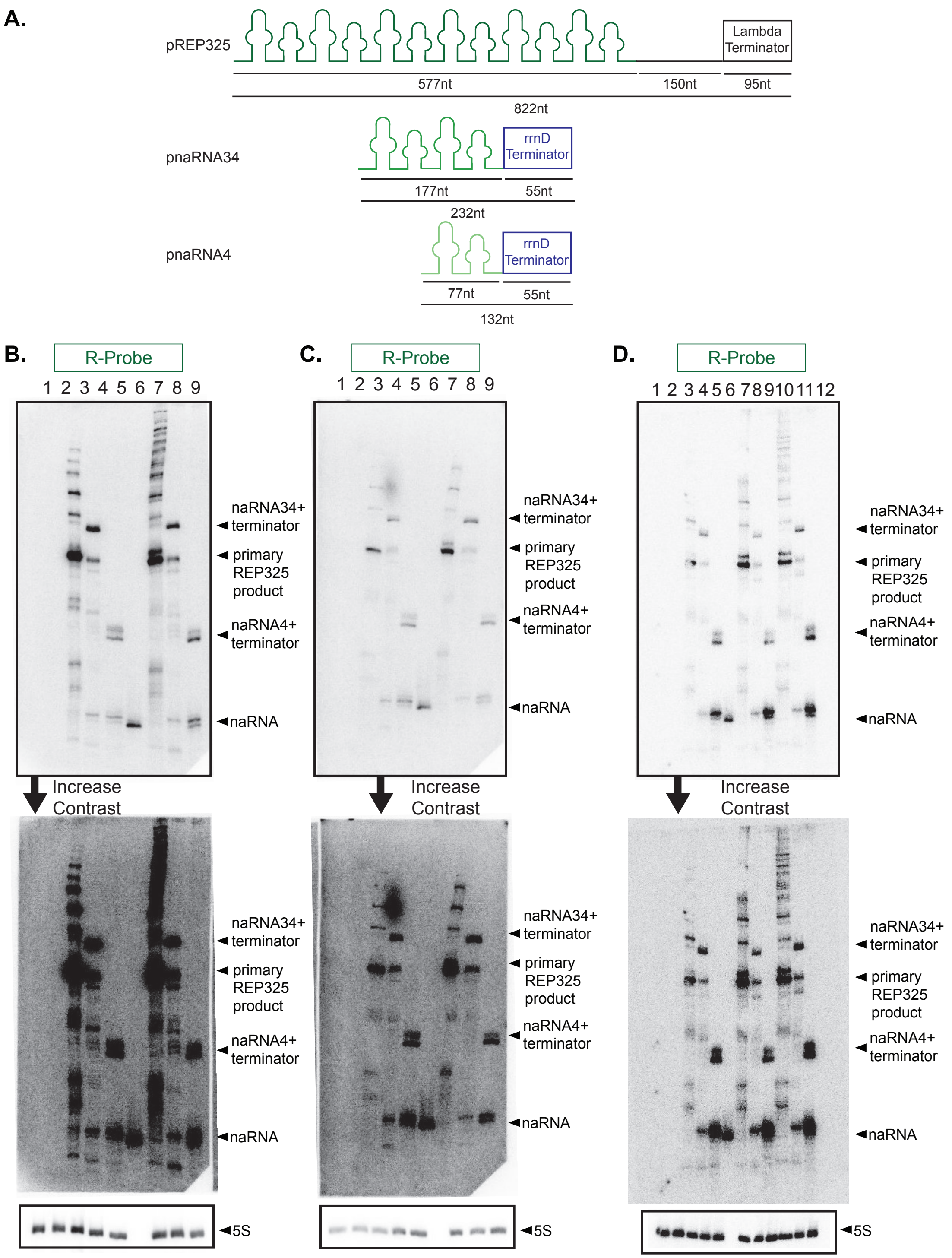

1 *wt*
2 *ΔREP325*
3 *wt*-pREP325
4 *wt*-pnaRNA34
5 *wt*-pnaRNA4
6 naRNA4 IVT
7 *wt*-pREP325 BCM
8 *wt*-pnaRNA34 BCM
9 *wt*-pnaRNA4 BCM

BCM treatment beginning at dilution

1 *wt*
2 *ΔREP325*
3 *wt*-pREP325
4 *wt*-pnaRNA34
5 *wt*-pnaRNA4
6 naRNA4 IVT
7 *wt*-pREP325 BCM
8 *wt*-pnaRNA34 BCM
9 *wt*-pnaRNA4 BCM

BCM treatment beginning at dilution

1 *wt*
2 *ΔREP325*
3 *wt*-pREP325
4 *wt*-pnaRNA34
5 *wt*-pnaRNA4
6 naRNA4 IVT
7 *wt*-pREP325 BCM - 30 minute treatment
8 *wt*-pnaRNA34 BCM - 30 minute treatment
9 *wt*-pnaRNA4 BCM - 30 minute treatment
10 *wt*-pREP325 BCM - treatment beginning at dilution
11 *wt*-pnaRNA34 BCM - treatment beginning at dilution
12 *wt*-pnaRNA4 BCM - treatment beginning at dilution

**Supplemental Figure 12. Replicates of *REP325* Northern blots A.** Schematic of plasmids used in Northern Blot for REP325 expression with respective lengths labeled **B.-D.** Biological replicates of Northern blots probing REP325 expression endogenously in *wt* and *ΔREP325* and overexpressed in *wt*-pREP325, *wt*-pnaRNA34, and *wt*-pnaRNA4 without and with BCM. 5µg (A.), 1.5µg (B.), and 5µg (C.) of total RNA was loaded for each lane. For IVT naRAN4, 0.2ng (A.), 0.05ng (B.), and 0.2ng (C.) was loaded for each lane

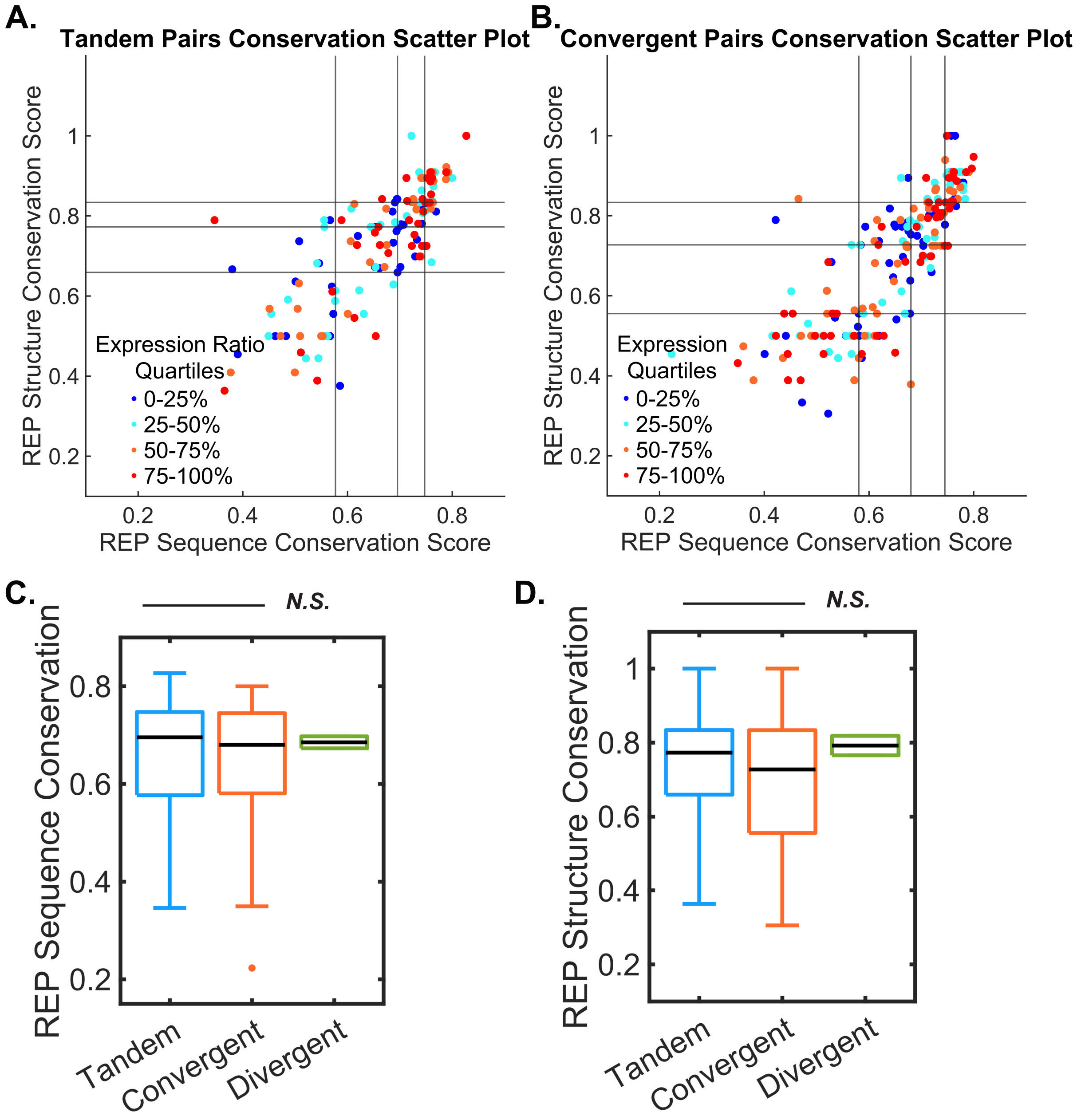

**Supplemental Figure 13. REP structure and sequence conservation in tandem and convergent genes A.** For tandem gene pairs, REP structure conservation vs REP sequence conservation with data quartiles indicated by vertical and horizontal lines, Spearman Correlation coefficient, r = 0.77, Data points are colored according to quartiles of upstream/downstream expression ratio. **B.** Same plot as A, but for convergent gene pairs with REPs. Spearman Correlation coefficient, r = 0.83. **C.** REP sequence conservation for REPs categorized by surrounding gene orientation. (Divergent only has 2 cases and cannot be statistically compared to the other conditions.) **D.** REP structure conservation for REPs categorized by surrounding gene orientation

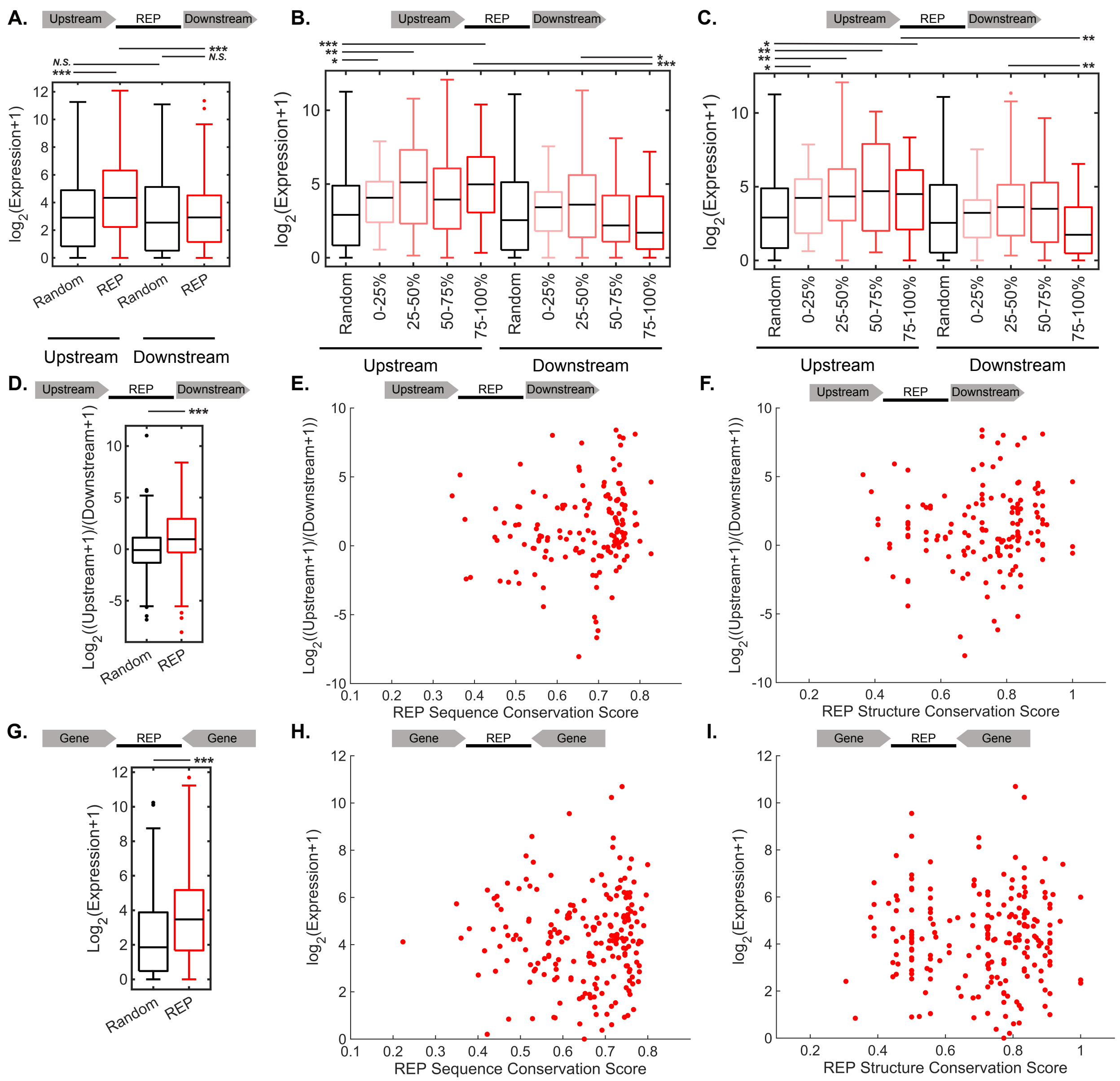

**Supplementary Figure 14 Genome-wide REP expression patterns A.** Absolute expression values of upstream and downstream genes without REPs and with REPs split **B.** Absolute expression values of upstream and downstream genes without REPs and with REPs split into quartiles by REP sequence conservation. **C.** Absolute expression values of upstream and downstream genes without REPs and with REPs split into quartiles by REP structure conservation. **D**. Expression ratio of upstream/downstream genes without REPs and with REPs, *p*-value = $1.2 \times 10^{-4}$ **E.** Correlation of expression ratios with REP sequence conservation for tandem genes. Spearman Corr = 0.202, *p*-value = 0.0144. **F.** Correlation of expression ratios with REP structure conservation for tandem genes. Spearman Corr = 0.12, *p*-value = 0.15 . **G.** Mean expression value of convergent gene pairs without REPs and with REPs, *p*-value = $1.48 \times 10^{-14}$ **H.** Correlation of expression values with REP sequence conservation for convergent genes. Spearman Corr = 0.044, *p*-value = 0.526 **I.** Correlation of expression values with REP structure conservation for convergent genes Spearman Corr = -0.01, *p*-value = 0.88

**Supplementary Information**

**Fisher's Exact Tests**

REP325 deletion seems to cause a slight stress response. To test if the stress response was related to changes in chromosome structure, we tested if the *ΔREP325* DE genes were HU-regulated, REP-adjacent, or supercoiling-sensitive.

If the loss of naRNA4-HU compaction is driving the transcriptional changes seen in *ΔREP325*, we would expect the differentially expressed genes to be also regulated by HU. Probing if the expression changes could be HU-dependent, we found that 24 of the 160 *ΔREP325* DE genes were HU-regulated, a significant overlap (**Table S11**, Fisher's exact t-test *p*-value = 0.0004). However, the 24 genes do not seem to be modulated by an HU-REP dependent mechanism. Only 5 of the 24 genes are REP-adjacent. The 24 genes could be loosely described as metabolism-related (*srlAE,gutM,sdhAC,aceF, malEP,manY*), acid stress-related (*gadWX*), and flagellar (*flgBCDEF*), all of which align with the *ΔREP325* DE genes more generally. This association could be attributed to both HU-regulated genes and *ΔREP325* DE genes being independently related to stress response pathways, rather than a direct HU-mediated mechanism. We found no significant overlap between HU-regulated genes(4) and REP-adjacent genes more generally (**Table S11**, Fisher's exact t-test *p*-value = 0.9356). These results suggest that HU is not driving the differences in *ΔREP325* and *wt* expression.

If REP-REP contacts were an important part of chromosome organization, deleting one REP could affect the expression of genes adjacent to other REPs due to chromosomal rearrangement and changes in accessibility. However, the *ΔREP325* DE genes did not have significant overlap with the REP-adjacent genes throughout the chromosome (24 genes overlap, Fisher's exact t-test *p*-value 0.82), suggesting that expression of distant REP-adjacent genes are not interdependent.

Lastly, since REPs have been proposed to regulate supercoiling via gyrase binding, deleting REP325 could change the chromosomal supercoiling distribution. To test this possibility, we examined genes sensitive to gyrase inhibitor, novobiocin, as a proxy for supercoiling-sensitive genes. The *ΔREP325* DE genes do not overlap with novobiocin-sensitive genes (**Table S11**, Fisher's exact test *p*-value = 0.22) (5).Overall, the transcriptional changes in *ΔREP325* do not seem to come from HU, REP-REP interactions, or supercoiling-mediated changes.

**Northern Sensitivity Estimate**

We did not observe any endogenous expression of REP RNA at lengths between 50-1000nt (**Fig. S11B-D**). The purified naRNA4 indicated that 0.2ng of RNA was easily detected. Assuming 90% of the total cellular RNA is ribosomal RNA, 0.2ng would be equivalent to 0.04% of the nonribosomal RNA loaded. By an alternative approximation based on the yield and starting concentration of cells in the total RNA prep, 0.2ng of naRNA4 would be equivalent to ~90 copies/cell. We do not have an absolute lower limit of detection, but the weakest band detected (the longest product from either pREP325 or pREP325-BCM) on each polyacrylamide-based northern blot was 55-fold, 72-fold and 123-fold lower in intensity than the corresponding IVT product.

**Table 1. Strains and Oligonucleotides**

| Strains | | | |
|---|---|---|---|
| Name | Genotype | Plasmid | Source |
| *wt* | F-, λ-, rph(+C, 667/687), dgcJ::IS1(+9bp, 991-999/1491) | | (3) |
| *ΔREP325* | F-, λ-, rph-1, *ΔREP325* | | This work |
| *wt ECGRC* | F-, λ-, rph-1 | | ECGRC |
| *yjdM* KO | F-, *Δ(araD-araB)567*, *ΔlacZ4787*(::rrnB-3), *λ*$^{-}$, *rph-1*, *Δ(rhaD-rhaB)568*, *ΔphnA733::kan*, *hsdR514* | | ECGRC |
| *wt*-pREP325 0mM IPTG | F-, λ-, rph(+C, 667/687), dgcJ::IS1(+9bp, 991-999/1491), | pREP325 | This work |
| *wt*-pnaRNA34 | F-, λ-, rph(+C, 667/687), dgcJ::IS1(+9bp, 991-999/1491), | pnaRNA34 | This work |
| *wt*-pnaRNA4 | F-, λ-, rph(+C, 667/687), dgcJ::IS1(+9bp, 991-999/1491), | pnaRNA4 | This work |
| wt-pVector | F-, λ-, rph(+C, 667/687), dgcJ::IS1(+9bp, 991-999/1491), | pVector | This work |
| wt-pLacI | F-, λ-, rph(+C, 667/687), dgcJ::IS1(+9bp, 991-999/1491), | pLacI | This work |
| *ΔREP325*-pREP325 | F-, λ-, rph-1, *ΔREP325* | pREP325 | This work |
| *ΔREP325*-pnaRNA34 | F-, λ-, rph-1, *ΔREP325* | pnaRNA34 | This work |
| *ΔREP325*-pnaRNA4 | F-, λ-, rph-1, *ΔREP325* | pnaRNA4 | This work |
| *ΔREP325*-pVector | F-, λ-, rph-1, *ΔREP325* | pVector | This work |
| *wt*-pM | F-, λ-, rph(+C, 667/687), dgcJ::IS1(+9bp, 991-999/1491), | pM | This work |
| *wt*-pMR | F-, λ-, rph(+C, 667/687), dgcJ::IS1(+9bp, 991-999/1491), | pMR | This work |
| *wt*-pMRN | F-, λ-, rph(+C, 667/687), dgcJ::IS1(+9bp, 991-999/1491), | pMRN | This work |
| *wt*-pMN | F-, λ-, rph(+C, 667/687), dgcJ::IS1(+9bp, 991-999/1491), | pMN | This work |
| *wt*-pN | F-, λ-, rph(+C, 667/687), dgcJ::IS1(+9bp, 991-999/1491), | pN | This work |

Coding mutation in *wt* was found by whole-genome sequencing. No coding mutations were found in *ΔREP325* by whole-genome sequencing.

| Oligonucleotides | | |
|---|---|---|
| Name | Sequence (5'-3') | Source |
| naRNA4 template | GGATCCTAATACGACTCACTATAGAGGCCGGATAAGG CGTTTACGCCGCATCCGGCAACGGTGCCGACTGCCT GATGCGACGCTTGCGCGTCTTATCAGGCC | IDT |
| naRNA4 template (reverse) | GGCCTGATAAGACGCGCAAGCGTCGCATCAGGCAGT CGGCACCGTTGCCGGATGCGGCGTAAACGCCTTATC CGGCCTCTATAGTGAGTCGTATTAGGATCC | IDT |
| Cruciform Top Strand – Cy3 | CCACCGCTCAACTCAACTGCTTTGCAGTTGAGTCCTT GCTAGG/3Cy3Sp/ | IDT |
| Cruciform Bottom Strand | CCTAGCAAGGGGCTGCTACCTTTGGTAGCAGCCTGA GCGGTGG | IDT |
| *REP325*-kan-ccdB-u1 | GCCAGTTAAGTGTAGTTCGAAATTTATACAGATG AGAGGCATAGGAACTTCAAGATCCCC | IDT |
| *REP325*-kan-ccdB-l1 | GAAATCTGAGTTTGTGA AAAAGAACTGATTGTATTGTGATTTATATTCCCCAGAA CATCA | IDT |
| *REP325*-del-ss-01 | TTTGTGAAAAAGAACTGATTGTATTGTGATGCCTCTCA T CTGTATAAATTTCGAACTACA | IDT |
| M-Probe | CGT TAT CTT CGT AGG TGT ATT CGG AGT TGC | IDT |
| N-Probe | GCGTTGGTTGTTGTTTGACGACATT | IDT |
| R-Probe | CGCAAGCGTCGCATCAGGCAGTCGGC | IDT |
| 5S-Probe | CACACTACCATCGGCGCTACGGCGTTTCACTTCTGAG TTCGGCATGGGGTC | IDT |

**Table 2. Raw EMSA Quantifications**

Attached in Supplementary Data.

**Table 3. Hill Equation Fits of EMSA Quantifications**

| | $K_d$ (nM) | Hill Coefficient, n |
|---|---|---|
| WT HU-naRNA4 | 32.35±3.51 | 0.96±0.01 |
| HU-P63A-naRNA4 | 41.24±5.01 | 1.01±0.12 |
| HU-triKA-naRNA4 | 89.62±9.17 | 1.49±0.24 |
| WT HU-polyU | 4333±135.7 | 2.71±0.25 |

Fitted Values ± Standard Error

**Table 4. All Nucleoid and Cell Dimension Quantifications**

Attached in Supplementary Data.

**Table 5. Nucleoid and Cell Dimension Comparisons**

| Metric | Condition | Measurement | Condition | Measurement | *p* |
|---|---|---|---|---|---|
| Cell Area | *wt* | 3.2±0.1 $\mu m^2$ | *ΔREP325* | 3.2±0.1 $\mu m^2$ | 0.66 |
| Nucleoid Area | *wt* | 1.89±0.06 $\mu m^2$ | *ΔREP325* | 2.0±0.1 $\mu m^2$ | 0.58 |
| Cell Area | *wt*-pnaRNA4 | 2.5±0.1 $\mu m^2$ | *wt*-pVector | 2.6±0.1 $\mu m^2$ | 0.21 |
| Nucleoid Area | *wt*-pnaRNA4 | 1.39±0.04 $\mu m^2$ | *wt*-pVector | 1.45±0.05 $\mu m^2$ | 0.60 |
| Cell Area | *wt*-pREP325 | 2.9±0.1 $\mu m^2$ | *wt*-pVector | 2.6±0.1 $\mu m^2$ | 0.02 |
| Cell Width | *wt*-pREP325 | 0.97±0.02 µm | *wt*-pVector | 0.92±0.01 µm | $2.6 \times 10^{-6}$ |
| Nucleoid Area | *wt*-pREP325 | 1.54±0.05 $\mu m^2$ | *wt*-pVector | 1.45±0.05 $\mu m^2$ | 0.28 |
| Cell Area | *ΔREP325* pnaRNA4 | 2.41±0.07 $\mu m^2$ | *ΔREP325* pVector | 2.49±0.07 $\mu m^2$ | 0.12 |
| Nucleoid Area | *ΔREP325* pnaRNA4 | 1.42±0.05 $\mu m^2$ | *ΔREP325* pVector | 1.47±0.06 $\mu m^2$ | 0.30 |
| Cell Area | ΔREP325 pREP325 | 2.75±0.08 $\mu m^2$ | *ΔREP325* pVector | 2.5±0.1 $\mu m^2$ | 0.05 |
| Nucleoid Area | ΔREP325 pREP325 | 1.48±0.05 $\mu m^2$ | *ΔREP325* pVector | 1.47±0.06 $\mu m^2$ | 0.74 |
| Cell Area | *wt*-pLacI | 2.9±0.1 $\mu m^2$ | *wt*-pVector | 2.6±0.1 $\mu m^2$ | 0.21 |
| Nucleoid Area | *wt*-pLacI | 1.22±0.05 $\mu m^2$ | *wt*-pVector | 1.45±0.05 $\mu m^2$ | 0.001 |
| Cells with Asymmetric Nucleoids | *wt*-pnaRNA4 | 0/146 | *wt*-pLacI | 12/148 | |
| Cells with Asymmetric Nucleoids | *wt*-pREP325 | 1/152 | *wt*-pLacI | 12/148 | |

*wt, ΔREP325, wt*-pLacI conditions are no IPTG. All other conditions are 1mM IPTG.
*p* - two-sided KS-test

**Table 6. Growth Curve Fit Parameters**

| Condition | Lag Phase Duration (Hours) | Doubling Time (Min) |
|---|---|---|
| *wt* 0mM IPTG | 2.3 ± 0.4 | 35 ± 2 |
| *wt*-pREP325 0mM IPTG | 2.2 ± 0.3 | 35 ± 1 |
| *wt*-pM 0mM IPTG | 6 ± 2 | 37 ± 2 |
| *wt*-pMR 0mM IPTG | 3 ± 1 | 36 ± 1 |
| *wt*-pMRN 0mM IPTG | 5.7 ± 0.7 | 37 ± 2 |
| *wt*-pMN 0mM IPTG | 2.5 ± 0.4 | 36 ± 2 |
| *wt*-pN 0mM IPTG | 2.9 ± 0.4 | 37 ± 2 |
| *wt*-pnaRNA34 0mM IPTG | 2.1 ± 0.4 | 36 ± 1 |
| *wt*-pnaRNA4 0mM IPTG | 2.6 ± 0.2 | 36 ± 1 |
| *wt*-pVector 0mM IPTG | 2.7 ± 0.3 | 35 ± 1 |
| *wt*-pLacI 0mM IPTG | 4.3 ± 0.2 | 40 ± 2 |
| *wt* 1mM IPTG | 2.3 ± 0.4 | 36 ± 1 |
| *wt*-pREP325 1mM IPTG | 2.6 ± 0.4 | 37 ± 1 |
| *wt*-pM 1mM IPTG | 6 ± 1 | 38 ± 2 |
| *wt*-pMR 1mM IPTG | 4 ± 2 | 36 ± 1 |
| *wt*-pMRN 1mM IPTG | 6.2 ± 0.9 | 37 ± 2 |
| *wt*-pMN 1mM IPTG | 2.6 ± 0.4 | 37 ± 2 |
| *wt*-pN 1mM IPTG | 2.7 ± 0.2 | 36 ± 1 |
| *wt*-pnaRNA34 1mM IPTG | 2.3 ± 0.5 | 36 ± 1 |
| *wt*-pnaRNA4 1mM IPTG | 2.7 ± 0.3 | 35 ± 1 |
| *wt*-pVector 1mM IPTG | 2.5 ± 0.4 | 36 ± 1 |
| *ΔREP325 0*mM IPTG | 2.1 ± 0.4 | 29 ± 2 |
| *ΔREP325*-pREP325 0mM IPTG | 2.1 ± 0.2 | 29 ± 2 |
| *ΔREP325*-pnaRNA34 0mM IPTG | 1.9 ± 0.4 | 29 ± 2 |
| *ΔREP325*-pnaRNA4 0mM IPTG | 2.1 ± 0.2 | 29 ± 2 |
| *ΔREP325*-pVector 0mM IPTG | 2.0 ± 0.2 | 28 ± 1 |
| *ΔREP325 1*mM IPTG | 1.9 ± 0.4 | 29 ± 2 |
| *ΔREP325*-pREP325 1mM IPTG | 2.1 ± 0.3 | 30 ± 3 |
| *ΔREP325*-pnaRNA34 1mM IPTG | 2.0 ± 0.3 | 29 ± 3 |
| *ΔREP325*-pnaRNA4 1mM IPTG | 2.0 ± 0.2 | 28.7 ± 0.8 |
| *ΔREP325*-pVector 1mM IPTG | 1.8 ± 0.4 | 28 ± 2 |
| *wt* ECGRC 0mM IPTG | 2.6 ± 0.3 | 36 ± 1 |
| *ΔyjdM* | 2.2 ± 0.1 | 30 ± 1 |

Mean ± Standard Deviation

**Table 7. Growth Curve Parameter Comparisons**

| Metric | Condition | Measurement | Condition | Measurement | *p* |
|---|---|---|---|---|---|
| $t_d$ | *wt* | 35±2 min | *ΔREP325* | 29±2 min | $1.3x10^{-60}$ |
| lag phase | *wt* | 2.3±0.4 hours | *ΔREP325* | 2.1± 0.4 hours | $1.3x10^{-4}$ |
| $t_d$ | *wt* | 35±2 min | *wt* ECGRC | 36 ± 1 min | 0.53 |
| lag phase | *wt* | 2.3±0.4 hours | *wt* ECGRC | 2.6 ± 0.3 hours | 0.07 |
| $t_d$ | *ΔREP325* pnaRNA4 | 28.7 ± 0.8 min | *ΔREP325* pVector | 28 ± 2 min | 0.52 |
| lag phase | *ΔREP325* pnaRNA4 | 2.0 ± 0.2 hours | *ΔREP325* pVector | 1.8 ± 0.4 hours | 0.28 |
| $t_d$ | ΔREP325 pREP325 | 30 ± 3 min | *ΔREP325* pVector | 28 ± 2 min | 0.24 |
| lag phase | ΔREP325 pREP325 | 2.1 ± 0.3 hours | *ΔREP325* pVector | 1.8 ± 0.4 hours | 0.10 |
| $t_d$ | *wt*-pnaRNA4 | 35 ± 1 min | *wt*-pVector | 36 ± 1 min | 0.04 |
| lag phase | *wt*-pnaRNA4 | 2.6 ± 0.2 hours | *wt*-pVector | 2.5 ± 0.4 hours | 0.36 |
| $t_d$ | *wt*-pREP325 | 37 ± 1 min | *wt*-pVector | 36 ± 1 min | 0.72 |
| lag phase | *wt*-pREP325 | 2.6 ± 0.4 hours | *wt*-pVector | 2.5 ± 0.4 hours | 0.56 |
| $t_d$ | *wt*-pLacI | 40 ± 2 min | *wt*-pVector | 36 ± 1 min | 0.02 |
| lag phase | *wt*-pLacI | 4.3 ± 0.2 hours | *wt*-pVector | 2.5 ± 0.4 hours | $1.1x10^{-7}$ |
| $t_d$ | *wt* | 35±2 min | *ΔyjdM* | 30 ± 1 min | $9.7x10^{-5}$ |
| lag phase | *wt* | 2.3±0.4 hours | *ΔyjdM* | 2.2 ± 0.1 hours | 0.03 |
| lag phase | *wt*-pM | 6 ± 1 hours | *wt*-pVector | 2.5 ± 0.4 hours | $9.6x10^{-8}$ |
| lag phase | *wt*-pMR | 4 ± 2 hours | *wt*-pVector | 2.5 ± 0.4 hours | 0.02 |
| lag phase | *wt*-pMRN | 6.2 ± 0.9 hours | *wt*-pVector | 2.5 ± 0.4 hours | $2.4x10^{-11}$ |
| lag phase | *wt*-pMN | 2.6 ± 0.4 hours | *wt*-pVector | 2.5 ± 0.4 hours | 0.72 |
| lag phase | *wt*-pN | 2.7 ± 0.2 hours | *wt*-pVector | 2.5 ± 0.4 hours | 0.31 |
| $t_d$ | *wt*-pM | 38 ± 2 min | *wt*-pVector | 36 ± 1 min | 0.12 |
| $t_d$ | *wt*-pMR | 36 ± 1 min | *wt*-pVector | 36 ± 1 min | 0.86 |
| $t_d$ | *wt*-pMRN | 37 ± 2 min | *wt*-pVector | 36 ± 1 min | 0.32 |
| $t_d$ | *wt*-pMN | 37 ± 2 min | *wt*-pVector | 36 ± 1 min | 0.74 |
| $t_d$ | *wt*-pN | 36 ± 1 min | *wt*-pVector | 36 ± 1 min | 0.11 |

*wt*, *ΔREP325, wt*-EGRC, *ΔyjdM, and wt*-pLacI are no IPTG. All other conditions are 1mM IPTG.
$t_d$ – doubling time, lag phase – lag phase duration, *p* – two-sided t-test

**Table 8. DeSeq2 Results for *ΔREP325 vs. wt***
Attached in Supplementary Data.

**Table 9. GO Term Analysis of *ΔREP325* DE Genes**
Attached in Supplementary Data.

**Table 10. HU-Regulated *ΔREP325* DE Genes**

| Gene | $log_2$(Fold Change) | adjusted *p*-value | REP-adjacent |
|---|---|---|---|
| flgC | -6.58 | 7.07E-189 | |
| flgF | -6.53 | 5.87E-109 | Yes |
| flgD | -6.32 | 1.82E-87 | |
| flgE | -6.06 | 4.12E-74 | |
| flgB | -5.65 | 2.84E-23 | |
| ydcI | -2.16 | 1.63E-11 | |
| yjdN | 3.37 | 1.80E-10 | Yes |
| gadX | 3.92 | 6.01E-07 | |
| ygiW | 2.04 | 4.16E-06 | |
| sdhC | -2.47 | 4.89E-06 | |
| fruB | 2.52 | 5.05E-05 | |
| srlA | 4.07 | 8.36E-05 | |
| gutM | 3.62 | 0.000785 | Yes |
| srlE | 2.93 | 0.00154 | |
| yfcG | 1.79 | 0.00162 | |
| sdhA | -2.27 | 0.00164 | |
| cysP | 2.04 | 0.00186 | |
| manY | 1.64 | 0.00196 | |
| gadW | 2.17 | 0.00289 | |
| malP | 1.81 | 0.00480 | |
| ybcW | 2.39 | 0.00505 | |
| aceF | -2.36 | 0.00606 | |
| dcuC | 2.80 | 0.00606 | Yes |
| malE | 1.95 | 0.00756 | Yes |

**Table 11. Fisher's Exact Comparisons**

| | | |
|---|---|---|
| | Not *ΔREP325* DE | *ΔREP325* DE |
| Not HU-Regulated | 4175 | 134 |
| HU-Regulated | 306 | 24 |
| *p*-value | Odds-Ratio | Confidence interval |
| 0.0004 | 2.44 | [1.56-3.83] |
| | Not *ΔREP325* DE | *ΔREP325* DE |
| Not REP-Adjacent | 4175 | 134 |
| REP-Adjacent | 656 | 24 |
| *p*-value | Odds-Ratio | Confidence interval |
| 0.82 | 1.04 | [0.67-1.63] |
| | Not *ΔREP325* DE | *ΔREP325* DE |
| Not Novobiocin-Sensitive | 4175 | 146 |
| Novobiocin-Sensitive | 243 | 12 |
| *p*-value | Odds-Ratio | Confidence interval |
| 0.93 | 1.02 | [0.74-1.39] |
| | Not REP-Adjacent | REP-Adjacent |
| Not HU-Regulated | 3678 | 631 |
| HU-Regulated | 281 | 49 |
| *p*-value | Odds-Ratio | Confidence interval |
| 0.94 | 1.02 | [0.74-1.39] |
| | Not REP-Adjacent | REP-Adjacent |
| Not Novobiocin-Sensitive | 3749 | 635 |
| Novobiocin-Sensitive | 210 | 45 |
| *p*-value | Odds-Ratio | Confidence interval |
| 0.17 | 1.27 | [0.74-1.39] |

**Table 12. DeSeq2 Results for *wt*-pM *vs. wt***

Attached in Supplementary Data.

**Table 13. GO Term Analysis of *wt*-pM DE Genes**

Attached in Supplementary Data.

**Table 14. Northern Quantification Values**

Attached in Supplementary Data.

**Table 15. REPs appearing in Rho-binding screen**

Attached in Supplementary Data.

**Table 16. REPs appearing as bidirectional terminators**

Attached in Supplementary Data.
